# The representation of priors and decisions in parietal cortex

**DOI:** 10.1101/2021.05.03.442155

**Authors:** Tom R. Marshall, Maria Ruesseler, Laurence T. Hunt, Jill X. O’Reilly

## Abstract

Animals actively sample their environment through orienting actions such as saccadic eye movements. Saccadic targets are selected based both on sensory evidence immediately preceding the saccade, and a ‘salience map’ or prior built up over multiple saccades. In the primate cortex, the selection of each individual saccade depends on competition between target-selective cells that ramp up their firing rate to saccade release. However it is less clear how a cross-saccade prior might be represented, either in neural firing or through an activity-silent mechanism such as modification of synaptic weights on sensory inputs. Here we present evidence from magnetoencephalography for two distinct processes underlying the selection of the current saccade, and the representation of the prior, in human parietal cortex. While the classic ramping decision process for each saccade was reflected in neural firing rates (measured in the event related field), a prior built up over multiple saccades was represented via modulation of the gain on sensory inputs from the preferred target, as evidenced by rapid frequency tagging. A cascade of computations over time (initial representation of the prior, followed by evidence accumulation and then updating) provides a mechanism by which a salience map may be built up across saccades in parietal cortex. It also provides insight into why evidence accumulation signals are present in parietal cortex, when inactivation of the region has been shown not to affect performance on single-trial tasks.

## Introduction

Far from being passive recipients of sensory information, both humans and animals actively sample the environment using their sensory organs. In rodents, active sampling processes include whisking and sniffing; in primates, the most important and best-studied process is the control of saccadic eye movements.

As an observer views a visual scene, several saccadic eye movements per second are generated in order to direct the eye’s small focal window to points of potential interest. However, each sample does not stand alone – instead information from multiple fixations is integrated to construct a ‘model’ of the full visual field^1,2^, and this model in turn acts as a ‘prior, influencing the selection of targets for future saccades^3^. Therefore, the process of active sampling may be viewed as an interplay between two concurrent processes with distinct characteristics:

Firstly, the brain must generate each individual sampling action. Since only one saccade is made at once, each individual saccade must be selected by a process of *competition* between representations of alternative possible saccadic targets^4^; the process is by necessity *winner-take-all* in that the eyes can only fixate one location at a time^5^ and must operate on a fast timescale, driven by dynamics that ensure a new saccade is made every few hundred milliseconds^6^.

Secondly, the brain must *integrate* the limited information gained from many individual saccades – not only to construct the visual scene, but relatedly, to inform the selection of targets for future saccades. Behavioural modelling suggests that the likely information value of future saccades is represented in a salience map^7,8^ that can be regarded as a Bayesian prior distribution over potential saccadic targets^3,9^. Far from having winner-take-all dynamics, this salience map must capture the *relative or probabilistic distribution* over all possible saccades. It therefore must integrate information across multiple previous saccades. The neural processes underlying the selection of individual saccades are, due to a combination of electrophysiological and modelling work, relatively well understood^10–13^, but the representation of the prior is less so; the latter is the focus of the current study.

The mechanism by which candidate saccadic targets ‘compete’ has been elucidated using evidence accumulation paradigms, designed to extend the saccade selection process over hundreds of milliseconds – most commonly random dot kinematograms (RDK) or ‘moving dots’ tasks^14^. Spatially-selective neurons in frontal and parietal cortical eye fields (FEF and LIP) track the accumulation of evidence in favour of a saccade to their response field^10–13^. Such evidence-tracking activity is consistent with a model in which each neuron’s firing rate at a given moment represents the log odds that its preferred location will be the target of the next saccade, effectively ramping up to target selection or down to target rejection^10,15^. The process can be described mathematically as a sequential probability ratio test^12^. A key element of this decision model is a winner-take-all competition between targets^16^. An influential biophysically-specified form of this model developed by Wang and colleagues describes how two pools of neurons compete with each other to determine the choice of saccadic target^17,18,19^. In the present study, we use the Wang model to identify MEG signatures of the saccade selection process for each individual saccades.

In contrast, the mechanisms by which a prior or model is built up across multiple saccades is less well understood. One mechanism by which a prior can be represented is via modulation of the baseline firing rate of target-selective neurons; This has been observed in monkey LIP. However, such an ‘active’ representation of the prior would be energetically costly over longer timescales^7,20^. A second candidate mechanism is that synaptic weights from neurons in the ‘input layer’ (visual cortex) to the layer representing the prior are adjusted to favour different sensory inputs^17^. The process of gain modulation, which is largely activity-silent, would be an energy-efficient way to store priors over longer timeframes.

To probe the gain modulation hypothesis, we developed a novel approach using human neuroimaging (magnetoencephalography – MEG), using the method of rapid frequency tagging. We reasoned that a change in synaptic weights should result in a change in gain for even irrelevant sensory stimuli co-located with favoured saccadic targets. In rapid frequency tagging, an irrelevant, invisible perceptual manipulation (high-frequency rhythmic flicker of the targets), known to produce strong increases in oscillatory power in sensory cortices^21,22^, is presented. These ‘tag’ oscillations propagate forward through the visual system, thus indirectly probing input gains to higher visual areas such as posterior parietal cortex. Importantly, two competing saccadic targets can be tagged with different frequencies so that the representation for the two targets can be precisely separated as it propagates forward through the visual system, even if populations of cells tuned to the different targets are partially or completely spatially intermixed. This approach is therefore sensitive to prior beliefs encoded via synaptic plasticity^17^ rather than neural firing rates.

Although cortical regions concerned with saccade selection exist in both frontal cortex (frontal eye field, FEF) and parietal cortex (lateral intraparietal area, LIP), this study focusses on the parietal cortex. The reason for this is mainly practical, as the visual frequency tag signal is attenuated as it propagates forward in cortex and is not detectable in the frontal lobe. However, we note that recent studies using cortical inactivation in rodents and monkeys suggest a key role of parietal saccade regions in construction of the prior. Rats with parietal inactivation showed a reduced influence of previous trials on interval judgements, and parietal neurons were shown to represent information about recent trials^23^. In contrast, inactivation of frontal eye fields (frontal orienting fields in the rat) affects performance on single trial target selection, but inactivation of parietal cortex does not^24,25^. Therefore the parietal cortex is a strong candidate for the neural substrate of the prior for saccadic selection.

## Results

We modified the classic random-dot kinematogram (RDK) task^26^ in two ways (fig1A) to generate distinct predictions about the neural signatures of both the competitive process of selecting a single saccade, and the integrative process of constructing and representing the prior.

**Figure 1:**
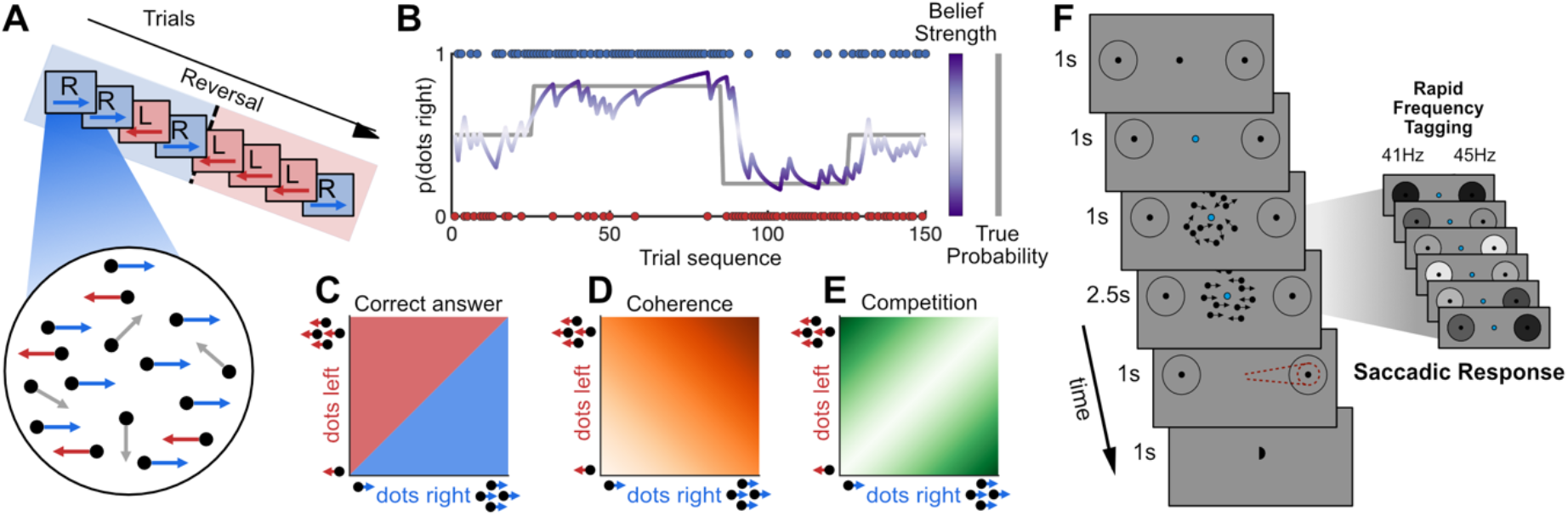
Adapted dot-motion task. A) Trial sequences were presented where certain motion directions were more common, with unpredictable ‘reversals’ (top), and stimuli within a trial varied along two orthogonal dimensions; number of dots moving left, and right, with all other dots moving randomly (bottom). B) the cross trial prior probability p(correct direction = right) varied across the experiment, with pseudo-random, unsignalled ‘blocks’ of trials in which the dominant direction was rightwards 20%, 50% or 80% or the time (grey line). Bayesian learning models were used to estimate direction and strength of beliefs and observer should have about the current trial based on previous trials (purple line). C) On a given trial, the correct response is given by the dominant motion direction. D,E) Additionally, 2-d stimulus space can also be parameterized as varying along two dimensions: **total coherence** (middle panel), or the total percentage of coherent motion to the left or right, and **competition** (right panel), the unsigned difference between proportion of coherent dots moving left and right. F) Structure of a single trial: A get-ready cue indicated dots were about to appear. All trials began with 1 second of random motion, which temporally separated stimulus onset from evidence onset, meaning we were able to distinguish visual evoked neural responses from evidence accumulation processes. This also provided a temporally extended foreperiod in which neural activity reflecting prior beliefs could be measured. Random motion was followed by a 2.5s evidence accumulation period, in which coherent motion was present. When the dots disappeared, participants had a 1s response interval to make a saccade in the direction of perceived dominant motion. Participants were given unambiguous feedback on the correct answer (small centrally-presented hemisphere on the correct side), so they could learn the cross-trial prior independently of the quality of evidence on each trial. During the 1s foreperiod and 2.5s period of evidence accumulation, the two potential saccadic targets were ‘tagged’ with high frequency flicker to selectively entrain neural oscillations.

In the classic RDK task participants observe mixtures of *randomly* and *coherently*-moving dots and accumulate evidence over hundreds of milliseconds to determine the direction of coherent motion, responding with a congruent saccade. We introduced longer term, *cross-trial integration* favouring left- or right-response, to drive the computation of a prior hypothesised to occur in parietal cortex. Additionally, we introduced *within-trial competition* between left- and right options so that the strength of evidence for a target could be dissociated from the direction of evidence (and therefore its concordance with the prior).

### Cross-trial integration (prior)

In classic RDK tasks, the dot direction on each trial is independent. Each option (left, right) is equally likely, and so subjects do not have to retain any information about the current trial after the trial ends. In our modified version, we introduced long-term correlations in the dot-motion direction. The probability that the next correct choice would be ‘right’ was not fixed at 50%, but took values of 20%, 50% or 80% for blocks of about 25 trials (changes in prior probability were un-signalled and occurred with a uniform hazard rate of 0.04 (fig1B, ‘true probability’ (solid grey line)). Since the dominant direction on previous trials could be useful for determining the direction on the current trial, participants could benefit from integrating information across trials to construct a *prior* over the dominant motion direction on the next trial (fig1B, ‘belief strength’ (purple trace)). We modelled this evidence integration process using a Bayesian ideal observer model (similar to ^27,28^), which captured local variations in prior probability, and learning delays. We used the expected value from the Bayesian ideal observer (see ‘Methods’ eq5) of p(rightward = correct), and modelled belief strength as |p(rightward) – 0.5|, meaning that it became strong when there was a high prior probability of either leftward or rightward saccade. This allowed us to test whether these prior beliefs were reflected in brain activity, either as an influence on the decision process itself or in the 1s foreperiod prior to evidence presentation on each trial (fig1F).

### Within-trial competition

In classic RDK stimuli a single set of coherent dots move to the left or right. In our modified version, all trials included some level of evidence for *both* choice options (left and right) concurrently, and participants reported the *dominant* motion direction (fig1C). This means that the level of the signal-to-noise or *total coherence* (total number of left and right dots, compared to random dots, fig1D). was manipulated orthogonally to *competition* (ratio of left to right dots, fig1E). Therefore, the strength of evidence on a given trial (the precision of the likelihood function in Bayesian terms) was dissociable from its direction, which could be compatible or incompatible with the prior. This allowed us to test (behaviourally and neurally) for evidence that the prior was used and precision-weighted against current evidence strength. This manipulation also allowed us to localize in time (and space) the single-saccade decision process, by testing for the hallmark of a competitive decision process namely that activity should depend upon the strength of evidence for the losing option as well as the evidence for the winning option^16^. In particular, we used Wang’s biophysical model of a drift diffusion-like competitive process to model decision making in parietal and frontal cortex^29^; this model makes precise predictions about the independent effects of competition and coherence on brain activity^30^.

#### Choice behaviour is influenced by the prior in accordance with Bayesian theory

Participants (n = 29, final analysis n = 26, for information on participant exclusions see ‘Methods’) performed 600 trials of a modified random-dot motion task with added within-trial competition and cross-trial integration, divided into 4 blocks with short breaks, while MEG was recorded. All participants also completed a practice session of 300 trials, outside the scanner, on a separate day.

We first confirmed that participants’ single-trial choice behaviour was influenced by both the total coherence (signal to noise) and the competition between left- and rightward motion, by fitting logistic regressions to participants’ saccade directions (left, right) as a function of percent coherent motion and proportion of coherent dots that moved right. As expected, there was an interaction such that participants were more likely to saccade rightwards when a greater proportion of the coherent dots moved rightwards and this effect increased as the total amount of coherent motion increased (t(25) = 9.74, p < 2*10^−10^, fig2A).

**Figure 2:**
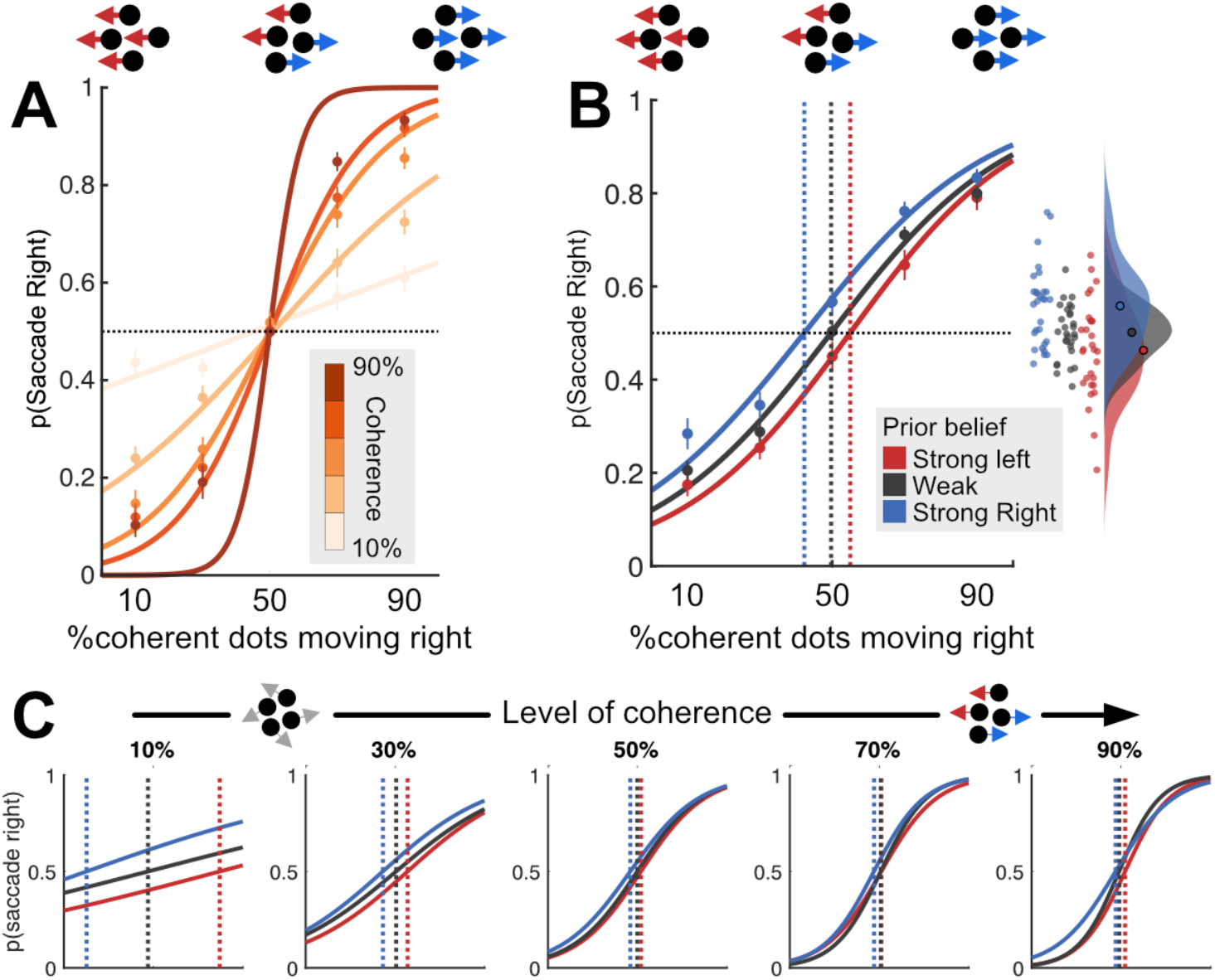
Within- and across-trial influences on behaviour. A) Increasing overall coherence (darker lines) produces a parametric performance improvement (steeper logistic curve). Dots indicate mean observed data, lines indicate logistic fits. B) Prior belief influences choice behaviour on current trial, leading to a shift in the point of subjective equality (dashed vertical lines). Inset: Raincloud plot showing values of p(Saccade right) for individual subjects on trials where 50% of dots moved right. C) Prior belief shifts point of subjective equality (dashed lines) most strongly when the least information is available on the current trial (coherence is lowest).

Next, we tested whether participants learned and used the across-trial regularities in the stimulus sequence (the prior).

We first confirmed that feedback from previous trials influenced participants’ decision on the current trial using lagged logistic regression (figS1), indicating that participants were indeed retaining information across at least two previous trials. To model participants’ prior beliefs, we used a Bayesian ideal observer model^27^ (see ‘Methods’ for full description) to compute, for every trial in a stimulus sequence, the *prior belief* that an ideal observer should have, based on the feedback observed on previous trials. The ‘ground truth’ or generative probability of rightwards motion was either high (80%), low (20%), or neutral (50%); these probabilities changed un-signalled about every 25 trials (see fig1 and ‘Methods’). The Bayesian ideal observer model estimated the probability that the dominant motion direction on the upcoming trial would be right or left, based on the feedback from previous trials. The advantage of this approach over using ‘ground truth’ prior probabilities is to capture local fluctuations in probabilities, and learning delays.

For visualisation (fig2B), we divided trials using a tertile split according to whether the Bayesian prior strongly favoured rightward or leftward motion, or neither (neutral prior). On ‘neutral prior’ trials, the point of subjective equality (PSE) closely matched the point of *objective* equality (mean 49.7% right motion). Strong priors in either direction biased the PSE by 10-15% (strong left prior; PSE 55% right motion, strong right prior PSE 42.3% right motion, fig2B, inset).

A multiple logistic regression confirmed that participants’ choice behaviour was influenced both by the proportion of dots moving right (t-test on regression coefficients across the group: t(25) = 14.79, p < 1*10^−14^) and by the prior based on previous trials (t(25) = 2.80, p = 0.0091). These two effects did not interact (t(25) = 0.45, p = 0.66). This is evidence that decision-relevant properties of the currently-viewed stimulus (the proportion of coherent dots that moved right), and the beliefs participants had developed based on previously-viewed stimuli, made independent contributions to their choice behaviour.

Because our bidirectional stimulus dissociated strength of evidence (total coherence) from direction of evidence (left vs right), we could test whether participants relied more upon the prior when evidence in the current trial was weak^31^ (low total coherence), due to precision weighting, as predicted by Bayesian theory^31^. To visualise this effect, we repeated the Bayesian ideal observer analysis but divided the trials according to the level of total coherent motion on the current trial (fig2C, using the same tertile splits as above). As predicted, prior belief biased current choice strongly when there was little decision-relevant motion, (at 10% coherence, PSE moved from ≈50% to 10%/90%, fig2C, left) but had little effect when a lot of motion was decision-relevant (at 90% coherence, PSE moved from ≈50% to 48%/52%, fig2C, right).

The effect shown in figure 2C was statistically confirmed using logistic regression. Percent total coherent motion had opposite effects on the influence of current evidence and prior belief on choice; when total coherent motion was high, the current evidence (proportion of coherent dots moving right) influenced behaviour more (Wilcoxon test on logistic regression coefficients for (coherence*prior) interaction across the group: Z = 4.7, p < 3*10^−6^, non-parametric test due to first-level outliers, see ‘Methods’) but prior belief influenced behaviour less (Z = −2.52, p = 0.012). This contrasts with the previous analysis where we found no interaction between the prior belief and the degree to which *competition* influenced choice behaviour. This confirms that participants selectively weighted the two sources of evidence available to them; up-weighting the impact of their prior belief – shaped by what they had seen *previously* - when little sensory evidence was *currently* available to guide their choice. This precision-weighting resembles the optimal Bayesian strategy for the task^31^.

#### Neural models and neuroimaging

Next, we turned to our MEG data for evidence of the neural mechanisms underlying the competitive process of selecting an individual saccade and the integrative process of forming a prior across many saccades.

##### Biophysical model of the neural mean field

We simulated the neural mean field (summed activity of neuronal pools representing the left- and right-targets) during the saccade selection process using an adaptation of the decision model previously described by Wang and colleagues^29,30^. Because Wang’s model describes the summed activity of all neurons in the decision-making population – the ‘neural mean field’ – it generates useful predictions about the population activity of a brain region measured by MEG. Indeed this model has been successfully used in a value-based choice task^30^ to predict the independent effects of total value (analogous to total coherence in our task) and value difference (analogous to competition in our task) on the event related field (ERF) as measured in MEG, and has previously been fit to brain activity in both parietal^29^ and frontal cortex^30,32,33^. We adapted a neural mean-field version of this model to make predictions about how neural responses would vary with competition, coherence and prior belief in our task.

Briefly, the model comprises two neuronal pools coding for different choice options, with strong recurrent excitation within a pool and strong inhibition between pools. The between-pool inhibition mediates a winner-take-all competition between options resulting in one pool reaching a high-firing attractor state and the other pool a quiescent attractor state. In the standard version of the model, inputs to the two pools are intended to be proportional to the strength of the input stimulus; in our case, number of dots moving left, and number of dots moving right.

To additionally model the effect of the prior as a driving input, we adapted the model to include *three* different input sources in different time windows (fig3A,B): The first input represented a participant’s prior belief about the upcoming stimulus (purple, fig3A, I_belief_, fig3B), the second reflected undifferentiated activity due to random dot motion during the 1-second ‘incoherent motion’ epoch (brown, fig3A), and the third reflected properties of the stimulus itself – i.e., the number of dots moving left and right – during the 2.5-second ‘coherent motion’ epoch (cyan, fig3A, I_dots_, fig 3B). The timing of the driving inputs reflects electrophysiological findings that the initial response of parietal neurons^34,35^ to target presentation, prior to evidence accumulation, weakly reflect the influence of prior beliefs on saccade selection^17,36^. Importantly, while the second input ‘general motion’, input was fixed across trials, the inputs corresponding to prior and to task-relevant motion were variable; figure 3B illustrates a trial where the visual stimulus more rightward than leftward motion (strong input to the ‘right’ pool), but the model expects leftward motion (strong input to ‘left’ pool).

**Figure 3:**
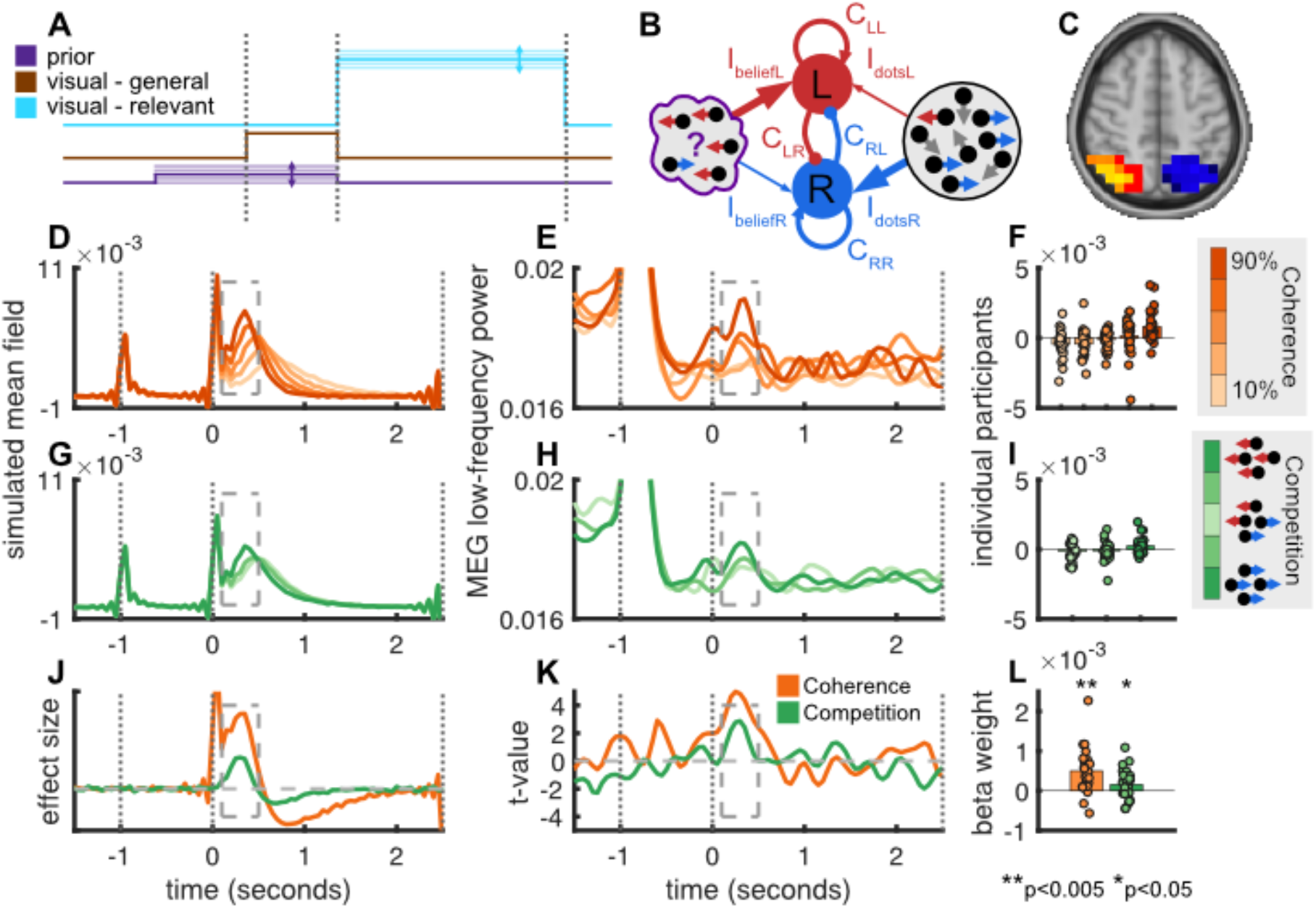
A) Input to neural mean-field model. Each node received weak variable input corresponding to prior knowledge (purple), weak fixed input corresponding to motion in all directions (yellow) and strong variable input corresponding to coherent motion (red). B) Properties of neural mean-field model on a representative trial. Each node contained recurrent excitatory connections and inhibitory connections to the other node, and received driving inputs corresponding to the strength of prior belief, and to the properties of the dot stimulus. C) Region-of-interest used to reconstruct activity in IPS (based on a combination of data and anatomy – see ‘Methods’). D) Neural mean-field model predicts a parametric increase in activity as a function of stimulus coherence. E); Low-frequency in IPS activity displays parametric modulation as a function of coherence. F) Time-averaged mean field from the period indicated by the box in E. Each dot is an individual participant. G) Neural mean-field model predicts a parametric increase in activity as a function of stimulus competition. H) Low-frequency in IPS activity displays parametric modulation as a function of competition. I) Time-averaged mean field from the period indicated by the box in H. Each dot is an individual participant. J) General linear model reveals effects of both coherence and competition in neural mean-field model in post-stimulus period. K,L) Significant effects of coherence and competition are observed in IPS in the same post-stimulus window.

To verify that this competitive model was a good fit to the underlying neural process, we tested for a key signature of a competitive selection process: namely that the decision variable depends on the strength of evidence for the *unselected* option, as well the selected option^16^. We were able to directly test for the influence of the unchosen option because in our modified random dots task, evidence for the chosen and unchosen dots direction was manipulated independently, allowing us to identify signals driven by competition independently of signal to noise.

The neural mean-field model generated three distinct predictions about the activity that should be observed in a competitive system. Firstly: Increasing total *coherence* (sum of dotsL and dotsR) should produce a parametric increase in neural activity following target onset (fig3D) in the time window 100-500ms following onset of coherent motion. Secondly: As *competition* decreases (i.e. greater absolute difference between dotsL and dotsR), there should be a parametric increase in neural activity following target onset (fig3G) in the same time window. Thirdly: Weakly increasing input due to prior knowledge in the prestimulus period (fig 3A, figS2A) should not significantly alter prestimulus activity, but rather bias the initial conditions of the competitive accumulation process such that activity evoked by the much stronger stimulus-driven input parametrically increased with prior strength, even though the prior input was no longer active, presumably due to the weak prior input biasing the state of the network before the stronger inputs began.

##### Event-related activity reflects the competitive process of selecting the current saccade

We next tested whether the pattern of results predicted by the neural mean-field model were observed in MEG data from the parietal cortex, which is known to track evidence accumulation. We transformed subjects’ MEG data to source space using LCMV beamforming^37^, and extracted time series from regions-of-interest (ROIs) in parietal cortex (fig3C, see ‘Methods’ for ROI definitions). We extracted the time-varying power in the low frequency range (2.8 – 8.4 Hz) as a proxy for the event related field or ERF, the magnetic field arising from local field potentials in cortex^38^; for further details see, ‘Methods / Parietal cortex and Frontal Eye Field low-frequency ROI analysis’. This analysis revealed two transient responses (fig3E,F,H,I); the first shortly after onset of incoherent motion, the second 100-500ms after onset of coherent (i.e., choice-relevant) motion (fig3E,H, dashed boxes).

Concordant with ramping activity observed in electrophysiological studies^10,12^, there was a clear parametric modulation of the low-frequency MEG signal 100-500ms after the onset of coherent motion as a function both of total coherence (fig3E,F) and stimulus competition (fig3H,I). As predicted, stronger evoked activity was observed with higher levels of total coherence; this is consistent with the observation that ramping evidence-accumulation signals in single-unit studies rise faster for higher coherence levels in standard dot motion tasks^12^. Stronger evoked activity was observed for *lower* levels of stimulus competition, indicating that the presence and strength of evidence conflicting with a decision affects the decision process – a hallmark of a competitive system. Statistically, a linear multiple regression with parameters coherence, competition and prior strength confirmed that the amplitude of the evoked response varied as a function of both coherence and competition; specifically, and as predicted by the neural mean-field model (fig3J) activity increased as a function of stimulus coherence (t(25) = 4.67, p = 9*10^−5^), and decreased as a function of competition (t(25) = 2.18, p = 0.039, fig3K,L). Both effects were tested in a time window 100-500ms after the onset of coherent motion (dashed boxes), defined based on the predictions of the biophysical model.

Since ramping activity is also observed in other brain regions such as FEF we repeated the analysis in this region (figS3). The pattern of results was qualitatively similar. To test whether there was any difference in timing of effects between FEF and parietal cortex, we used permutation testing (permuting the waveforms between ROIs within subjects), and found no significant difference in timing between regions (Coherence; p = 0.12, Competition; p = 0.92).

The qualitative correspondence of the MEG activity to a neural mean-field model of a competitive decision process suggests that parietal cortex could be engaged in resolving saccadic choices via competition by mutual inhibition, compatible with previous observations of evidence accumulation signals. However, it is worth noting that evidence from inactivation studies^24,39^ suggests that the activity in parietal cortex is not causally necessary for saccade selection (whereas activity in FEF is). The signal observed in parietal cortex may, we hypothesised, play a parallel role in integrating this information over a longer timeframe, into a cross-saccade prior.

##### Effects of the prior on the mean field

Contrary to our predictions, prior belief did not significantly modulate the low-frequency MEG signal in parietal cortex (t(25) = 0.80, p = 0.43, figS2). However, the absence of an effect of prior should perhaps be interpreted with caution, as for any null result; perhaps the effect was too subtle to be detected in the present paradigm. Furthermore, a relevant theoretical model^17^ predicts an interaction between signed evidence on the current trial and signed prior belief based on previous trials. Accordingly, we tested for this interaction, however no significant effects were observed in the low-frequency MEG signal in parietal cortex (left hemisphere; t(25) = −1.51, p = 0.14, right hemisphere; t(25) = 0.087, p = 0.93).

##### A prior integrating across multiple saccades is represented in parietal cortex via gain modulation

Although participants’ behaviour was clearly influenced by information on previous trials (figS1) consistent with computing a prior (fig2B), we were unable to detect activity corresponding to the prior in low frequency MEG signal (approximating the event related field, mainly driven by synchronised post-synaptic potentials).

We then turned to the possibility that the prior is represented via gain modulation. To probe for possible gain changes between visual inputs and the parietal cortex, we exploited the method of rapid frequency tagging^40^. During the time that moving dot stimuli were present on the screen the two saccade targets were rhythmically flickering at two different high frequencies (41 and 45Hz, fig1F) that were indistinguishable to observers. Flicker at such high frequencies is typically not perceived (they are for example close to the refresh rate of a standard 60Hz computer monitor – note that here we used a 1.44kHz projector to present stimuli). Indeed, no participant reported awareness that the stimuli were flickering. Flickering stimuli have previously been shown to produce detectable rhythmic activity in M/EEG signals^41^ including at higher frequencies above 40Hz^21,22^, presumably by producing synchronised neural activity in visual cortex that propagates through the visual streams^42,43^. Parietal neurons are known to increase their neuronal gain when an attentional or saccadic target is present in their receptive field^11,44–46^. Therefore, when one or other target is expected to be the saccadic target (for example due to a prior belief) we would expect the tag frequency for that target to propagate more effectively into parietal cortex; additionally we would expect the effect to be seen mainly in the contralateral hemisphere due to the lateralized representation of the visual field in occipital cortex.

### Validation of frequency tagging

In the MEG data we recovered clear spectral peaks at the 41Hz and 45Hz tag frequencies (fig4A) that were present while the tag stimulus was active. There was a clear lateralized effect such that the tag frequency presented at the right target was strongest in the left hemisphere of the brain, and vice versa.

**Figure 4:**
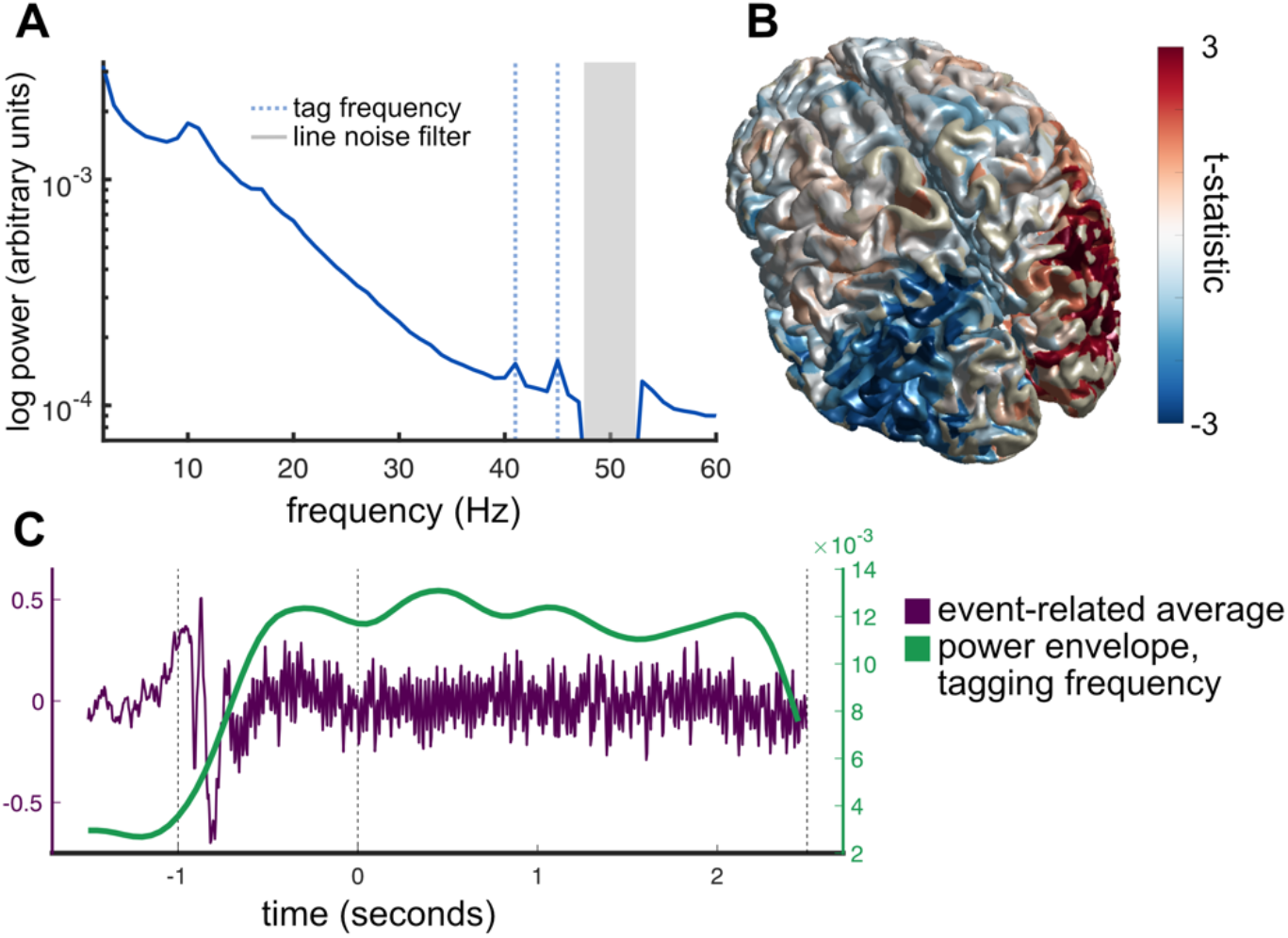
A) Average fourier spectrum of left and right parietal cortex ROIs displays clear peaks at the 41Hz and 45Hz tagging frequencies (dashed blue vertical lines). Note grey box indicates line noise filter. B) Unthresholded statistical map of differences between ‘power at left tag frequency’ and ‘power at right tag frequency’. Only parietal and lateral occipital regions showed lateralised differences in tag frequency power, presumably due to weaker SNR in frontal sources. C) Illustration of frequency tagging in a representative participant. Onset of moving dots and visual flicker at saccade targets produces detectable high-frequency activity in the timelocked-average MEG signal (grey trace). Sliding-window fourier analysis tracks the power envelope at this frequency (green trace).

The frequency tagging effect was strongest in parietal and occipital regions, propagating throughout posterior parietal cortex (fig4B), detectable from about 500ms after flicker onset (see ‘Methods’) and persisted during the entire stimulus period (fig4C). It should be noted that the frequency tagging signal could not be detected in frontal cortex including FEF. This was expected given that the effect is an entrained visual oscillation - the signal propagates forward from occipital cortex but only through a few synapses. Assuming some degree of post-synaptic ‘jitter’, more synapses between visual cortex and target region would necessarily lead to more jitter, and reduced signal, to the point where no tagging activity could be observed.

We defined ‘frequency tag activity’ for each parietal ROI (left, right) as the time-resolved power at the specific flicker frequency of the contralateral flickering saccade target, corrected for the main effects of flicker frequency and hemisphere. Notably, our parietal ROI overlapped substantially with two regions of interest thought to be homologous to eye-movement regions LIP and 7a in the macaque^47^; however the spatial resolution of MEG is not sufficient to say with certainty which intra-parietal sub-regions were the source. For full description of ROI definition see ‘Methods’.

For comparison, we also examined frequency tag activity in occipital cortex. If the prior is indeed represented as a gain modulation between the input layer (visual cortex) and the parietal cortex, we would expect to see task dependent modulation of the tag signal in parietal cortex, but not occipital cortex.

### The cross-saccade prior is represented prior to evidence accumulation

If the parietal cortex represents a prior expectation based on the integration of previous saccades, this should be in evidence in the foreperiod, when only incoherent motion was present. We defined a time window from 500ms after flicker onset (the point at which tag activity was first evident in parietal cortex – fig4) until the onset of coherent motion. We divided trials with a tertile-split into those on which the prior strongly favoured the contralateral target, strongly favoured the ipsilateral target, or was close to neutral.

Concordant with our hypothesis, we found that during the foreperiod, frequency tagging activity reflected the direction of the prior – activity was strongest on trials when the prior favoured the contralateral target (linear contrast comparing tertiles strong-contra, weak, and strong-ipsi: t(25) = 2.27, p = 0.028, fig5A,B). No such effect was observed in occipital cortex (t(25) = −0.223, p = 0.824, fig5C,D). To test for a parametric effect of prior strength on frequency tag strength, we conducted a linear regression of tag strength on prior strength (defined as the prior probability of contralateral motion on the upcoming trial, p(contra)) – this failed to reach significance (group t-test on regression coefficients, (t(25) = 1.50, p=0.073) suggesting that either the representation of the prior is largely categorical, or that we had insufficient sensitivity to detect a parametric effect.

**Figure 5:**
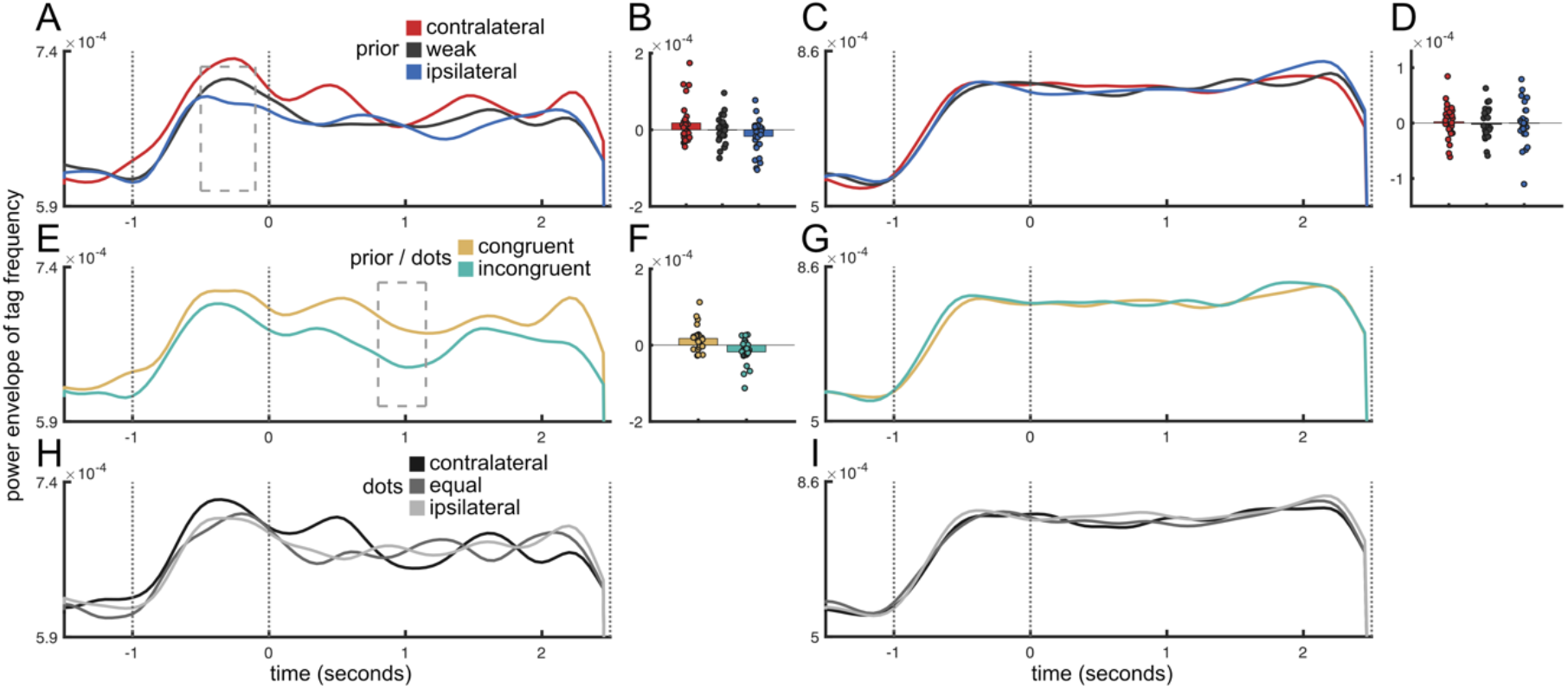
A) Time course of frequency tagging activity in parietal cortex as a function of prior belief (contralateral, weak, or ipsilateral to region). B) Direction of prior belief modulates parietal activity before onset of coherent motion. C,D) As A,B, but for occipital cortex. E) Time course of frequency tagging activity in parietal cortex as a function of congruence of prior belief and dominant direction of moving dots (congruent, incongruent). Dashed box indicates time-window where cluster values were maximal. F) Individual datapoints averaged across time-window denoted by cluster test. G) As E, but for occipital cortex. Note, no individual data points are plotted as nonparametric cluster permutation revealed no clusters. H) Time course of frequency tagging activity in parietal cortex as a function of dominant dot-motion direction. No significant dot-motion-related activity modulation was observed. I) As H, but for occipital cortex. No significant modulation was observed.

### Gain modulation in parietal cortex is sensitive to dot motion, **in the context of the prior**

While the frequency-tag data provided evidence of gain modulation relating to prior beliefs, it did not track evidence accumulation (coherence or competition on a single trial basis) in the same way as the ERF (figS4). This is compatible with a model in which the prior is represented as activity-silent synaptic plasticity^17^, or by tonic changes in baseline firing rate as in a bump attractor^48^ since tonic activity would likely not be detected in the band-passed MEG signal.

The gain modulation signal relating to prior beliefs was observed in the foreperiod of the task, during which the only lateralized effect is prior belief. If parietal cortex integrates incoming evidence with the prior belief to form a posterior, that will become the prior for the next trial (belief updating), we might expect to see a further gain-modulation signal representing the evidence, or decision, *relative to the prior*, reflecting this update process.

We coded trials as ‘congruent’ – i.e., the target favoured by the prior was also favoured by within-trial evidence – or ‘incongruent’. Indeed, stronger activity at the contralateral tagging frequency was observed on congruent than incongruent trials, during the evidence accumulation phase of the trial (fig5E,F; linear regression followed by cluster-based permutation test on regression coefficients, t_maxsum_ = 21.0023, cluster p = 0.0238 see ‘Methods’; no such effect was seen in occipital cortex (no cluster p < 0.05, fig5G)). Interestingly, the parietal effect was observed to be strongest in a later time window (800-1150ms after stimulus onset) than the decision-related activity. This suggests that, in the context of a paradigm in which information can be integrated across many saccades, evidence coding in parietal cortex represents the combination of this evidence with the prior, akin to the formation of a posterior distribution that could be used to guide selection of future saccades^27^.

At no point did we observe a main effect of the ultimate saccade direction in parietal or occipital cortex (no clusters p < 0.05, fig5H,I); this is perhaps unsurprising as the frequency tagging stimulus ended at the end of the coherent motion period, several hundred ms before the saccade was made.

### Lateralization of effects

It is notable that while the effects of prior belief observed in parietal cortex from frequency tagging (fig5) were lateralised, we were unable to detect lateralized effects in our analysis of the low-frequency, event-related responses (fig3, figS2). In that analysis, activity in parietal cortex *as a whole* resembled the neural mean-field model *as a whole* (for null analyses of lateralised effects of motion on the current trial and prior belief based on previous trials, in the low-frequency event-related MEG signal, see supplementary figures S5 and S6 respectively). Frequency tagging capitalizes on the exquisite spectral specificity of the MEG signal – due to their differing peak frequencies, signals originating from the left and right target could be very clearly distinguished and contrasted (see the spectral peaks in figure 4A).

### No effect of task variables on parietal tagging signal in occipital cortex

To determine whether our findings were specific to parietal cortex, we repeated the analysis of prior belief, prior/stimulus congruence, and dot direction (fig4) on the frequency tagging signals from the occipital cortex (figS8). No effects of any task variables were observed. This supports the interpretation that the prior is represented as a change in gain between occipital and parietal regions, such that tags associated with both saccadic targets are represented equally in occipital cortex, but visual information from receptive fields containing the target favoured by the prior propagates preferentially to parietal cortex. Additionally, the absence of task modulation gives us confidence that we are recording separate signals that behave differently. Even though we used beamforming to separate contributions from distinct neural sources, it is possible, due to field spread, that – rather than representing independent signals from parietal and occipital cortex respectively – the frequency tagging channels in fact represent two different measurements of the same underlying neural source. This control analysis makes that possibility unlikely.

#### Time course of neural effects suggests a sequential interplay between the prior and evidence accumulation processes

An advantage of MEG over other human neuroimaging methods is its high temporal resolution. This allowed us to examine the sequence of effects concerning the selection of individual saccades and their integration into a prior (fig6).

**Figure 6:**
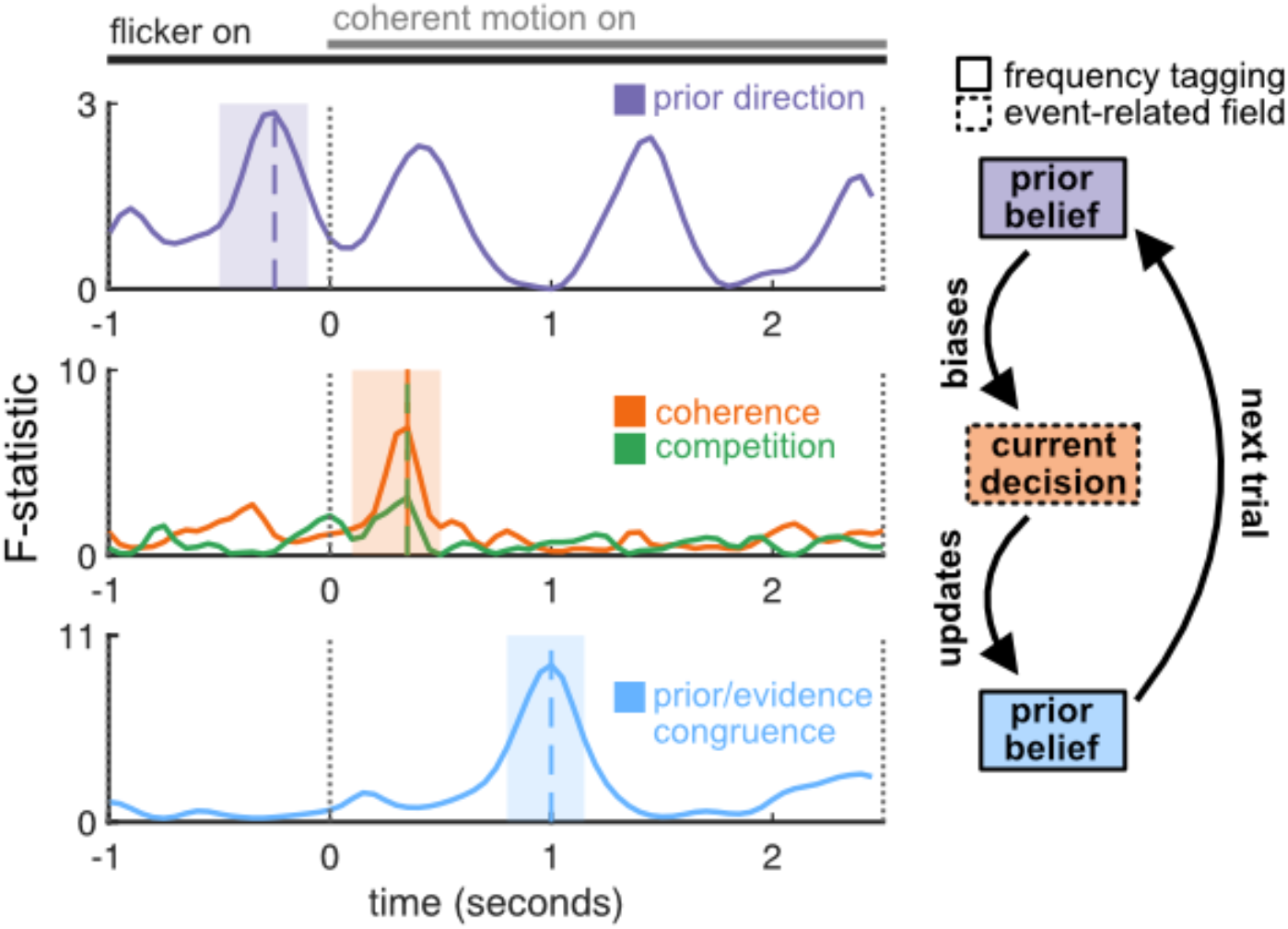
Summary of neural effects. Temporal cascade of computations. Time courses of significant statistical effects; repeated measures ANOVAs comparing contralateral, weak, and ipsilateral prior belief (top row), all levels of stimulus coherence (middle row, orange) and stimulus competition (middle row, green) and comparing ‘evidence (dots) congruent with belief’ trials with ‘evidence (dots) incongruent with belief’ trials (bottom row). Shaded boxes indicate preselected analysis windows (top and middle rows, see fig3E,F,H,I and 4C,D) and window delineated by cluster permutation test (bottom row, see fig4E,F). Coloured vertical dashed lines indicate maxima of the respective F-statistics.

To directly compare the sequence of events we ran a series of ANOVAs plotting the effect sizes for each of the previously described factors affecting parietal activity, as a function of time. A clear sequence of events emerged: initially the prior is represented via gain modulation (frequency tag effect), then the evolution of event-related activity reflects the evidence accumulation process as captured in a competitive mean field model (event-related field), and finally the interaction of evidence and prior is again reflected in gain modulation (frequency tag effect). Before the onset of coherent motion (interval [-1 0]), differences in activity across trials must relate to the prior. In this ‘foreperiod’ time window, we indeed observed a lateralized effect in parietal cortex as a function of the participants’ prior belief. Then, shortly after stimulus onset we observed parametric effects of stimulus coherence and stimulus competition predicted by the neural mean-field model and likely the signature of the choice process itself. Finally, having resolved competition between choice options, we observed a later effect which related to congruence between the participants’ initial prior (i.e., a prediction about the upcoming stimulus direction) and their eventual decision; namely, a strong suppression of activity when the stimulus is incongruent with the prior belief. This likely reflects the integration of evidence from the current trial with the prior belief – perhaps reflecting a *belief update* process whereby information about the current trial is used to modify the state of parietal cortex to guide future saccades.

The above is consistent with a model in which event-related neural activity field reflects the evidence accumulation process, but a prior integrating over multiple previous saccades and their outcomes is also represented by the modulation on input gains in parietal cortex^17^. This prior is represented even outside the time period in which evidence for different saccades is being weighed. The representation of the prior that is present before each saccade (in the foreperiod in our experimental task) serves to bias the saccade selection process, influencing choice behaviour (fig2B,C). During evidence accumulation, competitive dynamics lead to the selection of one or other saccadic target. Once the saccade selection process is resolved (but before the saccade is physically executed), an ‘update’ signal is observed in parietal cortex, perhaps reflecting the integration of the new evidence into the prior belief to form a posterior.

## Discussion

Optimal exploration of the visual environment through eye movements requires the brain to perform two distinct computations: selecting among *currently* competing saccadic targets, and integrating information across saccades to construct a Bayesian prior in order to plan *future* saccades.

Using whole-brain imaging at high temporal resolution with MEG, combined with computational modelling, we demonstrated a temporal cascade of computations relating to both these processes in parietal cortex. Notably our observations are consistent with distinct mechanisms for the processes. Competition between left- and right-targets is resolved by mutual inhibition between two pools of neurons driven by left- and rightwards evidence. This process produces characteristic signals relating to the strength of evidence both for the chosen and unchosen option, which can be detected in both FEF and parietal cortex. In addition, prior beliefs are represented through gain modulation in parietal cortex, in advance of evidence presentation; after the competitive process is resolved, this signal reflects the congruence between current evidence and prior beliefs.

Neurally, representing each type of task-relevant information – currently available and previously learned – requires computations that operate on different intrinsic timescales, and display different neural dynamics. Resolution of competition between two currently-available choice options requires a fast, winner-take-all neural system that rapidly converges to one of two stable attractor states. In contrast, representation of previously-learned knowledge must maintain a broadly similar state over a long timescale and incrementally change as new datapoints are incorporated. It is therefore perhaps unsurprising that the two computations are implemented by different neural mechanisms.

Our results provide evidence for a specific mechanism by which parietal cortex has been proposed to represent prior beliefs over longer timescales^17^, namely selective modulation of the gain between visual and parietal cortex. However, theory suggests that representations sustained over inactive periods or delays may be more efficiently represented by non-spiking mechanisms^20^ than active representation such as enhanced baseline firing rates. Physiologically, gain modulations could manifest through short term synaptic plasticity on the timescale of seconds. This is reminiscent of the proposal that short term synaptic plasticity may act as an activity-silent store for working memories^17^.; the activity pattern associated with working memory is later reactivated as a bump attractor^48,49^. Indeed a prior over saccadic targets could be viewed, in terms of timescale, as a form of non-cognitive working memory.

The method of rapid frequency tagging, while limited to regions of cortex within a few synapses of V1, can be used as a tool to probe activity silent representations in human cortex which cannot be accessed by studying standard stimulus-evoked fields^50^, and could be considered a temporally extended form of ‘pinging’^51^. This may be particularly germane to the study of representations constructed and maintained over slow timescales, particularly those spanning multiple task trials or events. An additional strength of the method is that, by exploiting the spectral precision of the MEG signal, the representation of stimuli ‘tagged’ with different frequencies can be disentangled, even when neurons tuned to those stimuli are intermixed in cortical tissue.

## Methods

### Behavioural task

Ethical approval for the study was granted by the local ethics board (Medical Sciences Interdivisional Research Ethics Committee, R53476/RE004). Participants (N=34) performed a variant of the classic dot-motion task. 1 second after a preparatory cue a field of 100 dots covering 4.2 degrees of visual angle (on-screen width 8.8cm, screen distance 120cm) was presented at fixation for 3.5 seconds. For the first second all dots moved randomly (uniform distribution over motion directions). After one second some dots changed direction and moved either to the right or left. There were five levels of overall coherence (sum of left-moving and right-moving dots): 10%, 30%, 50%, 70% and 90%, and three levels of competition: Coherently-moving dots either moved 90% or 70% in the direction of dominant motion, or 50% in each direction (i.e., no correct answer). Two saccade targets subtending 4 degrees of visual angle were concurrently presented at 6.8 degrees eccentricity, 0.8 degrees below the horizontal midline. Participants were instructed to maintain fixation until the dots disappeared, then to make a saccade to the saccade target on the side of dominant motion as quickly as possible. After 1s feedback (a semicircle at fixation) was given indicating the correct side.

Across trials the correct side was drawn from a generative distribution with p(right) either 20%, 50%, or 80%. The generative distribution changed randomly across a block with a fixed probability of 4%, i.e., a switch approximately every 25 trials. On trials with no correct answer (i.e., where equal numbers of dots moved left and right) feedback was drawn from the same generative distribution as the dots, however a programming error led to the feedback in this condition being the reverse of what was intended. This error affected approximately 8% of all trials.

All participants were trained on the task in a separate session outside the MEG for 300 trials (approximately 40 minutes). 5 participants were excluded due to low accuracy in the practice session and 29 participants completed the MEG session. During MEG acquisition data were recorded in four runs of max 20 minutes, each consisting of 150 trials. Total task time was approximately 80 minutes.

### Power calculation

Sample size was determined by simulation. Based on the z-scores and sample size from the MEG analysis in^30^, we calculated the effect size d as the z-score divided by the square root of the sample size. We then simulated a population of 10,000 virtual participants with this effect size, randomly sampled subsets of 10-60 participants from this population, calculated the sub-sample z-score, and computed statistical power for each sub-sample size as the proportion of z-scores for that sample size that exceeded 1.65 (equivalent to p < 0.05 in a one-tailed test). Simulation results indicated statistical power of 80.5% at our final sample size of N=26.

### Neural Models

#### Neural mean-field model

To model decision dynamics in Frontal Eye Field and parietal cortex we implemented a mean-field reduction of a spiking network model^18^. Full details of the biophysical parameters of the model are given in^30^. Briefly, the model consists of two connected neuronal pools each coding for one motion option (left, right), with within-pool excitatory connections and across-pool inhibition. Each unit receives noisy background inputs simulating endogenous cortical noise, plus three task-related inputs: Firstly; a weak (0 to 1.6Hz) input to one pool in the pre-stimulus period, simulating an initial bias in the decision process due to parietal input. Secondly; a weak (5Hz) input to both pools during the incoherent motion period, capturing the presence of low levels of motion in both decision-relevant directions. Thirdly; a strong input (ranging from 10.1Hz to 18.1Hz) to each pool in the coherent motion period proportional to the number of dots in each motion direction.

We focused on the synaptic inputs (I_1_, I_2_) to facilitate comparability with the MEG data, since MEG is known to be primarily sensitive to postsynaptic potentials (Hamalainen 1993). For optimal comparison with the MEG data which was bandpass-filtered and transformed to the frequency-domain, removing the DC component, we used the temporal derivative of the signal from the mean-field model.

#### Bayesian learning model

Because the modified dots task had temporal structure (dominant motion direction on trial *j* could be predicted, but not perfectly, from trials *1… t−1*), performance could be facilitated by tracking the true generative distribution of dominant motion directions, including when it changed. To model participants’ learning strategies we used a Bayesian ideal observer model; a ‘virtual participant’ that was fed the sequences of feedback given to the human participants and constructed, via Bayesian inference, a belief about the generative distribution on the upcoming trial.

#### Model details

On each trial, the prior probability that the dominant motion direction would be ‘right’ followed a Bernoulli distribution with parameter *q_t_*, i.e. the prior probability that the correct answer would be ‘right’ on trial *t* was *q_t_*.

The prior distribution over *q_t_* was initiated as a uniform on the range (0,1) on the first trial, and thereafter obtained from the posterior over *q_t−1_*, based on the outcomes *x*_1:*t*−1_, combined with a uniform ‘leak’; the posterior and ‘leak’ distributions were weighted by a factor H representing the true hazard rate:

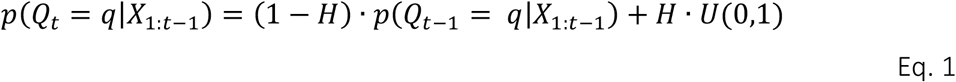

Where:

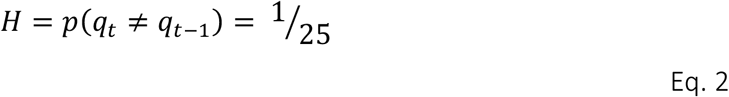

i.e. it was assumed that participants know approximately the true value of *H* following extensive pre-training.

The posterior *p*(*Q_t_* = *q*|*x*_1:*t*_) was obtained iteratively from the prior *p*(*Q_t_* = *q*|*x*_1:*t*−1_) and the likelihood:

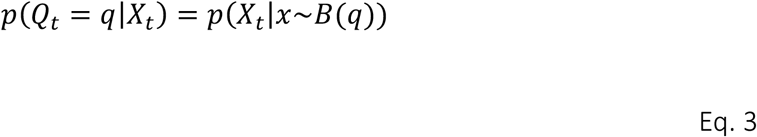

… using Bayes’ theorem:

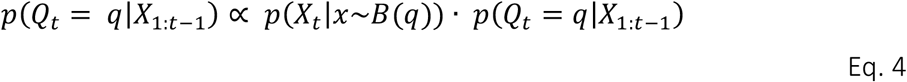

where the posterior over *q_t_* was normalized to integrate to 1.

Where a scalar value for the ‘prior’ is used in data analysis, this is the expected value of *q_t_* based on the prior distribution

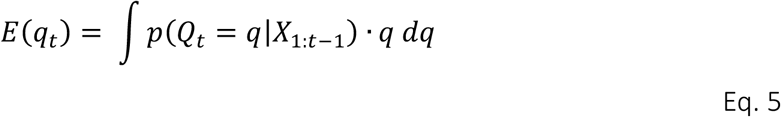

### Analysis of saccade data / behavioural data

Custom matlab code extracted saccade direction (left, right, or no saccade) based on the eyetracker data. Due to a technical error one participant’s eyetracker data were over-written and saccade information was reconstructed from the horizontal EOG.

To analyse saccade data we fit logistic regression models. Firstly, we asked whether the probability of saccading to the right on trial *t* depended on the proportion of coherent dots moving right on trial *t* (%R = 10,30,50,70 or 90%), the total coherence on trial *t* (coh = 10,30,50,70 or 90%), and the interaction of these (%R-mean(%R) x coh) (fig1A). Secondly, we asked whether the probability of saccading to the right on trial *t* depended on the proportion of dots moving right on trial *t* (%R = 10,30,50,70 or 90%), the participants’ prior belief about the dots on trial *t* formed from observing feedback on trials *1,2… t-1 E*(*q_t_*) as defined in Eq 5 above), and the interaction of these (fig1B).

Thirdly, we asked whether the observed effects of proportion dots moving right and prior belief on saccade direction were altered as a function of the level of stimulus coherence. To test this we fit a first-level logistic regression with evidence (%R = 10,30,50,70 or 90%), and prior belief *E*(*q_t_*) at each level of total coherence (coh = 10,30,50,70 or 90%). We then computed linear contrasts over the effects for evidence and prior belief across the five levels of total coherence. Due to the presence of outliers at the first level (generalized extreme studentized deviate many-outlier procedure^52^) we used nonparametric Wilcoxon signed rank tests at the second level to test against the null hypothesis of zero median.

### MRI acquisition

To enable localisation of cortical source generators of the MEG signal, a high-resolution structural MRI was acquired for each participant using a Siemens 3T PRISMA MRI scanner with voxel resolution of 1 × 1 × 1 mm^3^ on a 232 × 256 × 192 grid. The anatomical MRI scan included the face and nose to improve co-registration with the MEG data (see ‘MEG processing and analysis’). MRIs could not be acquired for 3 participants due to drop-out and screening contraindications. All imaging analysis was therefore conducted on the remaining 26 datasets.

### MEG acquisition

Data were recorded at 1.2KHz with an Elekta Neuromag VectorView 306 MEG system with 102 magnetometers and 102 pairs of orthogonal planar gradiometers. Head position indicator (HPI) coils were placed at four locations on the head to record head position relative to the MEG sensors at the start of each run. Head landmarks (pre-auricular points and nasion) and 200 points on the scalp, face and nose were digitized using a Polhemus Isotrack II system. EEG Electrodes were placed above and below the left eye and on the temples to record horizonal and vertical EOG, and on each wrist to record ECG.

### Rapid Frequency Tagging

During presentation of the dot-motion stimulus, the two saccade targets flickered rapidly at 41Hz and 45Hz (counter-balanced across subjects). The targets flickered sinusoidally between black and white (greyscale) at a refresh rate of 1.44KHz. To achieve this high rate of presentation we used a PROPixx DLP LED projector (VPIxx Technologies Inc., Saint-Bruno-de-Montarville, Canada). In post-experiment debriefing no participant reported awareness of the flicker.

### MEG processing and analysis

MEG analysis was performed using fieldtrip^53^, OSL (https://github.com/OHBA-analysis/osl-core), and custom MATLAB scripts.

Data were first Maxwell filtered using the MaxFilter program (Elekta Instrument AB, Stockholm, Sweden); Maxfilter is a method for separating parts of the recorded MEG signal that arise from external noise and neuronal activity respectively. MRI and MEG data were co-registered using RHINO (Registration of Headshapes Including Nose in OSL). Data were downsampled to 200Hz, bandpass filtered between 1 and 80Hz and bandstop filtered around the line-noise frequency of 50Hz. Trials containing outlier values were automatically detected and removed using function osl_detect_artefacts with default settings, and Independent Component Analysis was used to automatically remove additional artifacts associated with eye-blinks, ECG, and line noise.

A single-shell forward model was constructed from each participant’s anatomical MRI. Sensor data were projected onto an 8mm grid using an LCMV vector beamformer^37,54^ carried out on each 20-minute MEG run separately. The grid was constructed using a template (MNI152) brain, then each participants’ anatomical MRI was warped to the template brain and the inverse warp applied to the grid. This ensures comparability of source reconstructions across participants. Because MaxFilter considerably reduces the dimensionality of the data – to approximately 64 – the data covariance matrix was reduced to 50 dimensions using PCA. Eigenvalue decomposition of magnetometer and gradiometer channels was performed in order to normalise each sensor type to ensure that both sensor types contributed equally to the covariance matrix calculation^55^.

### Parietal cortex and Frontal Eye Field low-frequency ROI analysis

The parietal ROI was defined with respect to both anatomy (the Harvard Oxford cortical atlas) and function (band-limited power at the specific frequency-tag frequencies) – see ‘Frequency tagging analysis’. This was used to construct ‘virtual channels’ for left and right FEF based on a symmetric orthogonalization method^56^.

Frontal Eye Field was defined with reference to a probabilistic atlas of human visual and oculomotor cortical regions^57^.

A limitation of beamforming is that the polarity of the recovered signal cannot be disambiguated; source activity is inherently *sign-ambiguous*. This means that conventional event-related averaging in the time domain is problematic. Accordingly, we performed time-frequency analysis and extracted time-varying power in the low frequency range (2.8 – 8.4 Hz) as a proxy for the event related field or ERF, the magnetic field arising from local field potentials in cortex^38^, as power is not sign-ambiguous but rather always positive. Oscillatory power was then computed via time-frequency analysis using a 500ms sliding window multiplied with a Hanning taper, at 0.8Hz frequency resolution. We then averaged across the low-frequency 2.8-8.4Hz band as a proxy for evoked activity, resulting in a ROI-specific single-trial time series. We focussed on low frequencies rather than higher frequencies because the network model is dominated by low-frequency responses and does not exhibit higher frequency oscillations^30^.

Inspection of the MEG data revealed two evoked responses with comparable durations; one 100-500ms following the onset of the incoherent motion, and one 100-500ms following the onset of coherent motion. To visualise the effects of task variables on these evoked responses we took the average low-frequency power across each time window for each level of coherence (10%, 30%, 50%, 70%, 90% of dots moved either left or right, fig1E,F), competition (50%, 70% or 90% of coherently-moving dots moved in the same direction, fig1H,I), and prior strength (unsigned value output from Bayesian model *abs*(*E*(*q_t_*) − 0.5), figS2). Because the latter is a continuous variable we performed a tertile split into ‘weak’ (p(right) close to 0.5), ‘medium’, and ‘strong’ (p(right) close to 0 or 1) values for visualisation purposes.

To determine the effect of stimulus variables (coherence competition) and prior belief strength on low-frequency power in the FEF and parietal cotex we applied linear regressions with these three predictor variables in two ways: Firstly, for visualisation, at every time point (fig3E,H,K, figS3A,C,E). Secondly, for statistical purposes, over the average power values in a preselected box from 100-500ms after coherent motion onset (fig3F,I,L, figS3B,D,F), based on the results from the neural mean field model (fig3D,E,J).

### Frequency tagging analysis

To analyse the effect of the rapidly flickering saccade targets on posterior brain regions we used a combination of anatomical and data-driven selection criteria to focus on the brain regions that produced the strongest tagging response. We selected a 3s time-window from 800ms before the onset of coherent motion to 2.2s seconds afterwards - i.e., almost the entire period the targets were flickering - and calculated the fourier transform at all voxels for all participants and averaged across all artifact-free trials. We then compared power at the left-target frequency (41Hz for half the participants and 45Hz for the other half) with power at the right-target frequency, which revealed strong effects of tagging in voxels in parietal and occipital regions. We then used an anatomical atlas to create weighted maps defined by the conjunction of statistically significant differences between tag-frequency power and the parietal cortex anatomical label. The left parietal ROI consisted of all voxels in left parietal cortex that showed significantly greater power at the left-hemisphere tagging frequency than the right, and the reverse was true for the right parietal ROI. We multiplied these weight maps with the source-space data to create ‘virtual channels’ in left and right parietal cortex. We then performed time-frequency analysis on each virtual channel at the relevant tag frequency (41Hz or 45Hz, depending on the flicker rate of the contralateral frequency tag), using a longer 1000ms sliding window to increase frequency resolution. This produced time-resolved estimates of power at the relevant tag frequency in the left and right parietal ROIs.

For the analysis of prior belief (fig4C,D) we expected effects prior to coherent motion onset and therefore focused on the 1-second epoch of incoherent motion, focusing on a time window 500-100ms before coherent motion onset due to low SNR in the first part of the incoherent motion epoch as frequency tagging activity ramped up (fig4A).

We performed a tertile split on the prior, but – as we expected *hemispherically lateralised* effects due to the retinotopic organisation of parietal cortex – we used the signed prior, splitting into ‘strong left’, ‘weak’, and ‘strong right’. We then averaged tag power values in the selected time window, at each level of prior and compared these using a 1 × 3 ANOVA with linear contrast. We also conducted a linear regression using the parametric prediction of the prior probability of dots moving right, *E*(*q_t_*) as per Eq. 5, as explanatory variable, in the same time window.

Because we did not have a strong a priori prediction about the time window in which parietal activity might be driven by belief-decision congruence we used a nonparametric cluster-based permutation test^58^ over all time points to determine whether there was a difference between the congruent and incongruent conditions. We again performed a tertile split on the prior; strong left, weak, strong right, and a three-way split on the dot directions; mostly left motion (10% or 30% right), equal motion (50% right), or mostly right motion (70% or 90% right). We then defined trials where the dominant motion direction and prior agreed as ‘congruent’, and trials where they were opposite as ‘incongruent’, and performed the cluster-based permutation test on the participant conditional means.

### Time course analysis

To illustrate how the computational components of active sampling evolve in time we calculated, for every time point in the task epoch (from the onset of incoherent motion until the cessation of coherent motion) repeated measures ANOVAs comparing relevant task variables. We compared, respectively; frequency tagging activity in parietal cortex as a function of prior strength (strong contralateral, weak, strong ipsilateral), low-frequency activity in FEF and parietal cortex as a function of stimulus properties (coherence; 10%, 30%, 50%, 70%, 90%, competition; low, medium, high), and parietal frequency tagging activity as a function of prior/stimulus congruence (congruent, incongruent). To illustrate the temporal cascade of events we found the time point at which each F-statistic was maximal.

## Acknowledgements

The authors would like to thank Sven Braeutigam for technical assistance with data collection, and Eelke Spaak for helpful comments on an earlier version of the manuscript.

JOR is supported by a Career Development Fellowship from the Medical Research Council (MR/L019639/1). MR is supported by a PhD studentship from the Wellcome Trust (109064/Z/15/Z). LTH is supported by a Sir Henry Dale Fellowship from the Royal Society and the Wellcome Trust (208789/Z/17/Z). The Wellcome Centre for Integrative Neuroimaging is supported by core funding from the Wellcome Trust (203139/Z/16/Z).

This research was funded in whole, or in part, by the Wellcome Trust. For the purpose of Open Access, the author has applied a CC-BY public copyright licence to any Author Accepted Manuscript version arising from this submission.

## Data availability statement

The data underlying this manuscript are available upon request to the corresponding author. Data will be deposited in a public repository upon publication of a revised manuscript.

## Code availability statement

The code used to generate these results is available upon request to the corresponding author. Analysis code will be deposited in a public repository upon publication of a revised manuscript.

## Supplementary Information

### Lagged logistic regression reveals integration kernels over previous trials

**Figure S1:**
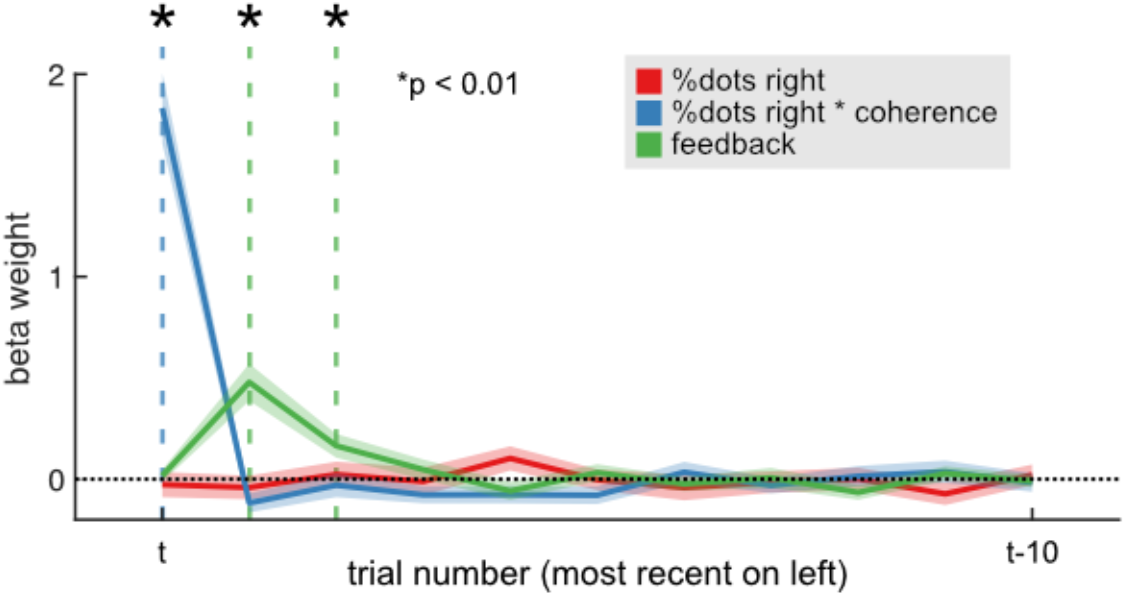
Lagged logistic regression of current and previous task variables on current choice. Error bars indicate standard error of the mean across participants.

To determine whether participants indeed integrated information from previous trials we performed a logistic regression with dependent variable; saccade direction (left, right), and independent variables; proportion of coherent dots moving right, interaction of proportion coherent dots moving right with coherence (effectively the absolute number of dots moving right) and post-trial feedback. To determine whether task parameters on *previous* trials influenced choice on the *current* trial we included the above regressors at ‘lags’ ranging from zero (i.e., the task parameters on the current trial *t*) to 10 (parameters on trial *t-10*).

Lagged multiple regression revealed that absolute number of right-moving dots on the current trial strongly predicted choice on the current trial (t(25) = 10.51, p = 3.2e-11), but on previous trials (lags > 0) did not predict current choice. In contrast, feedback on trial *t* did not predict choice on trial *t* (t(25) = 0.56, p = 0.6); this was entirely expected since feedback *followed* choice. However, feedback on trial *t-1* (t(25) = 5.2, p = 2e-5) and trial *t-2* (t(25) = 2.8, p = 0.009) did predict choice on trial *t*. Proportion of dots right on current or previous trials did not predict current choice beyond the other regressors (all t < 1.8).

The above strongly indicates that participants’ choice on trial *t* was influenced by the stimulus properties on trial *t*, and the history of post-trial feedback on previous trials *t-1* and *t-2*.

### No evidence that prior belief strength modulates low-frequency activity in frontal or parietal cortex

**Fig S2:**
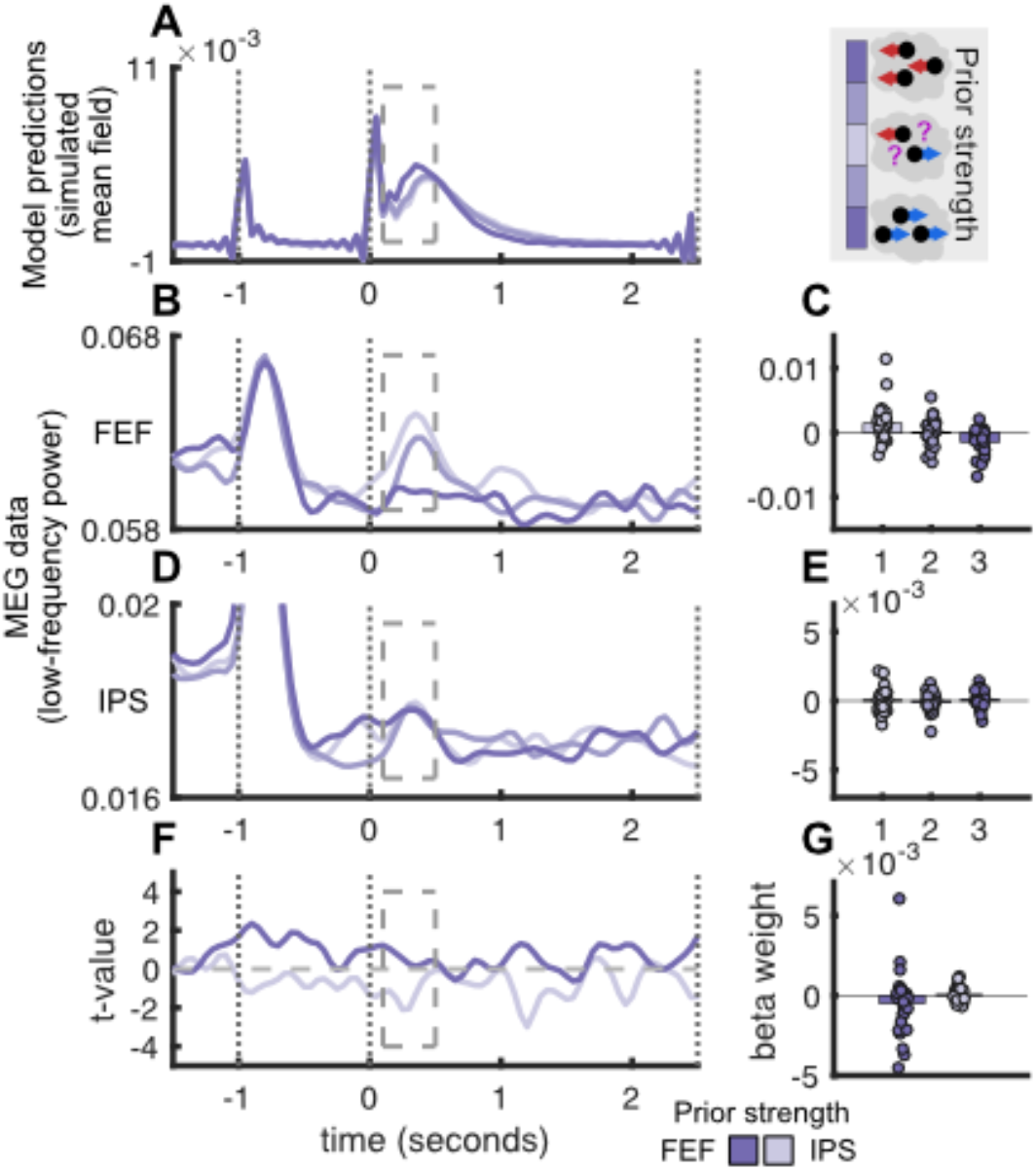
Signals related to strength of prior belief. All plotting conventions as fig3, main text. A) Neural mean-field model predicts a parametric increase in activity as a function of strength of prior belief. B,C: FEF low-frequency power. D,E: Parietal cortex low-frequency power. F,G: Results from general linear model.

Our neural mean field model (fig3A, main text) operationalised prior belief as a weak input in the prestimulus period. However, rather than showing strong changes in *prestimulus* signal, the model rather exhibited changes in *stimulus-evoked* activity, presumably by biasing the prestimulus state. A strong prestimulus input produced the strongest stimulus-evoked activity (figS2A). However, in contrast to this prediction we did not observe significant changes in stimulus-evoked activity in the MEG data. In FEF we observed a non-significant effect in the opposite direction; stimulus-evoked activity was strongest on trials where participants had a *weak* prior belief close to 50/50 (general linear model, t(25) = −1.22, p = 0.23, figS2B,C). In parietal cortex no trend was observed in either direction (t(25) = 0.8, p = 0.43, figS2D,E).

### Stimulus properties parametrically modulate low-frequency activity in Frontal Eye Field similarly to Parietal cortex

**Fig S3:**
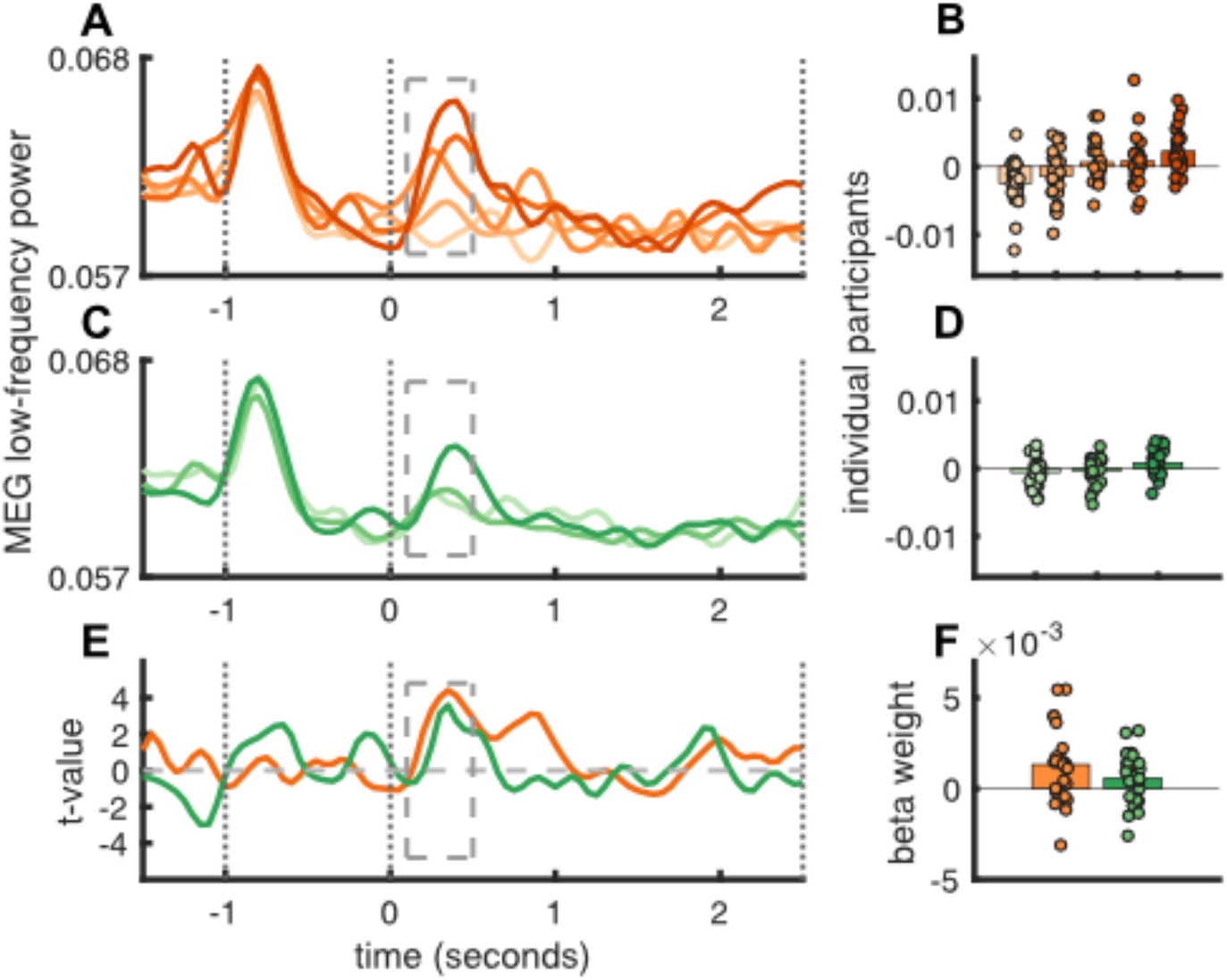
As fig3, but plotting the effects of stimulus coherence (A,B) and competition (C,D)) on low-frequency activity in the Frontal Eye Field. E,F: GLM analysis confirms significant effects of both variables in the 100-500ms following stimulus onset.

As ramping activity has been observed in neurons in FEF as well as parietal cortex, we repeated our analysis of stimulus variables on ERF activity in the FEF (for FEF ROI definition see ‘methods’). As for parietal cortex, a multiple linear regression with parameters coherence, competition and prior strength confirmed that evoked response amplitude varied as a function of both coherence (t(25) = 3.42, p = 0.0021), and competition (t(25) = 2.13, p = 0.043) in the 100-500ms window after coherent motion onset (dashed boxes), defined based on the biophysical model (fig3).

### Effects of stimulus variables (coherence, competition) on frequency-tagging activity in parietal cortex

For comparison with the analysis of stimulus-driven effects (fig3) we applied the same model to the frequency tagging data from parietal cortex, pooled across left and right hemisphere ROIs. As we were able to observe lateralized effects of the prior in parietal cortex (fig4), we also tested for a lateralized effect of evidence, i.e., whether ‘more dots moving towards the contralateral side’ produced a stronger tag response than ‘more dots moving towards the ipsilateral side.

Visually inspection suggested no strong effects of either coherence (figS4A,B) or competition (figS4C,D) on or signed/lateralized competition (figS4E,F) in the frequency tagging signal. To confirm this we fit a general linear model with these three effects to the frequency tagging data on single trials, then comparing the beta-weights from the first level model to zero. No significant effects were observed (coherence, t(25) = −0.37, p = 0.72; competition, t(25) = - 0.93, p = 0.36; lateralized competition, t(25) = 1.02, p = 0.32).

**Figure S4:**
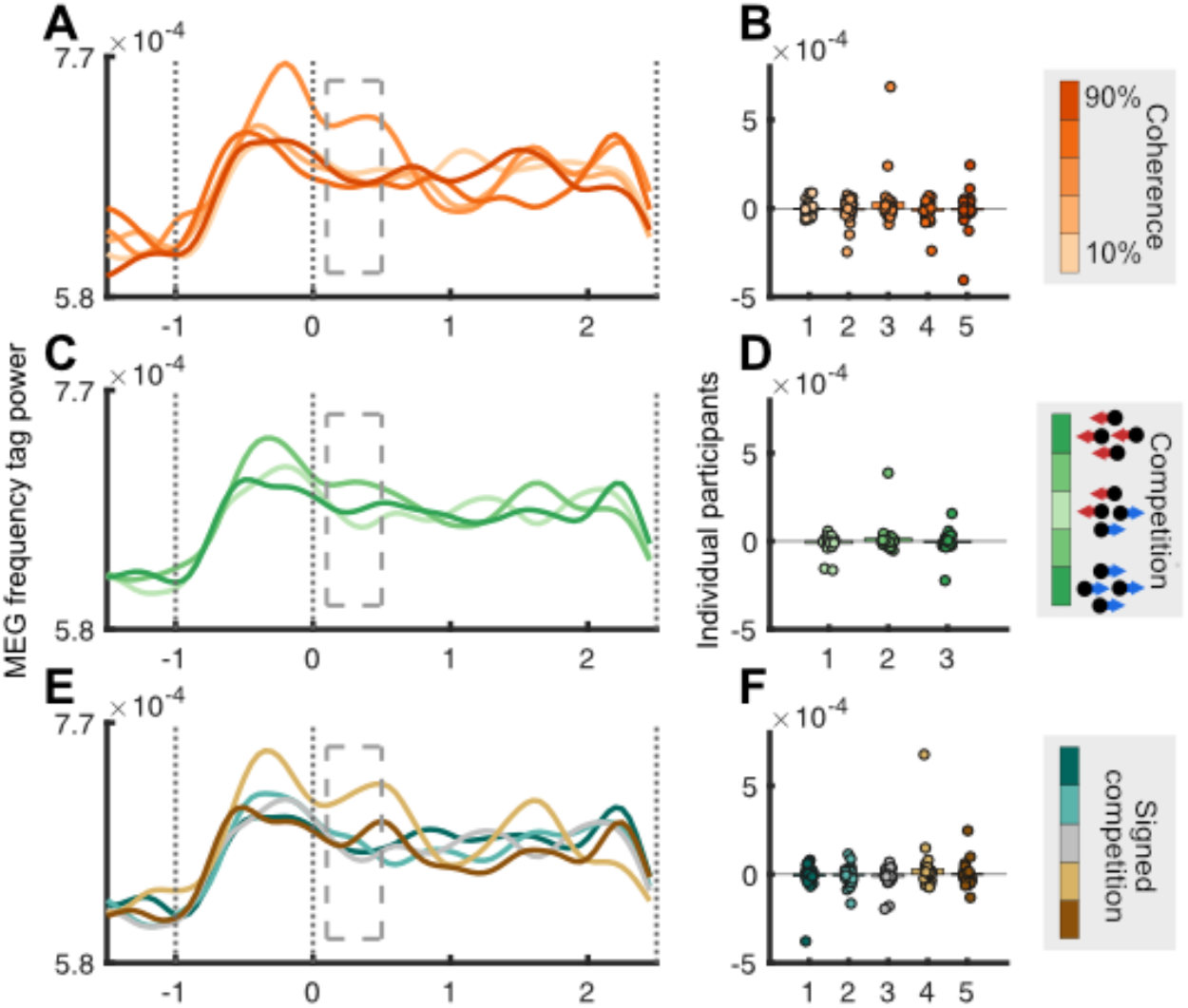
Analysis of parietal cortex frequency tagging data analogous to low-frequencies (fig3) A) Parietal frequency tagging time series as a function of stimulus coherence. B) Averaged across 100-500ms time window, dashed box in A. C,D) As A,B, but as a function of stimulus competition. E,F) As A,B, but as a function of ‘signed competition’, i.e., proportion of coherent motion moving contralateral to region of interest.

### Ability to detect lateralised task processing in Frontal Eye Fields and Parietal Cortex

Frontal Eye Field and LIP neurons predominantly code for saccades to the contralateral hemispace^59,60^. Therefore we might have expected the left- and right-hemisphere regions of interest to show lateralized effects in addition to the overall evidence accumulation signals reported in figure 3.

To obtain lateralized predictions, we looked at the inputs to individual pools in the neural mean-field model, rather than the mean field produced by summing all inputs (see main text). In the model our prediction was confirmed; as contralateral coherent motion increased, the simulated evoked response grew larger (figS5A,B). However, we did not observe a comparable effect in the FEF MEG data (fig S5C-F). Observation of the evoked-response window suggested a U-shaped pattern, where strong evidence for *either* motion direction (10%, 90% conditions) produced a large evoked reponse, whereas high competition (50% condition) produced the weakest evoked response.

Fitting a general linear model with % coherent dots moving right as a regressor confirmed an absence of evidence for left and right FEF acting as individual evidence accumulators. No significant linear effect was observed in either the left FEF (t(25) = −0.61, p = 0.55) or right FEF (t(25) = −0.96, p = 0.35). In fact qualitatively the left FEF data appear to show a quadratic effect entirely consistent with the main effect of competition (fig3); trials with the largest number of dots moving in one direction produced the strongest evoked responses, and 50/50 trials with maximum competition produced the weakest.

In parietal cortex (figS5G-J) a different pattern was observed: Here a linear effect of proportion coherent dots moving right was observed in the left (t(25) = 3.32 p = 0.003) but not the right hemisphere (t(25) = −0.86 p = 0.40).

The absence of evidence for our two predictions in FEF could be due to a lack of hemispherically-lateralised decision processing in FEF. It may be the case that neurons accumulating evidence for both choice options are present in both hemispheres. However, an alternative possibility is that we are unable to fully disentangle signals from left and right FEF via beamforming^37^. Similarly, in parietal cortex neurons responding to left- and right visual fields are not completely segregated, with the left hemisphere showing a more lateralized response than the right^61^, perhaps consistent with the fact that the lateralized signal in parietal cortex was strong in the left hemisphere. However, again the limitations of beamforming may partly explain the difficulty in detecting lateralized responses.

**Figure S5:**
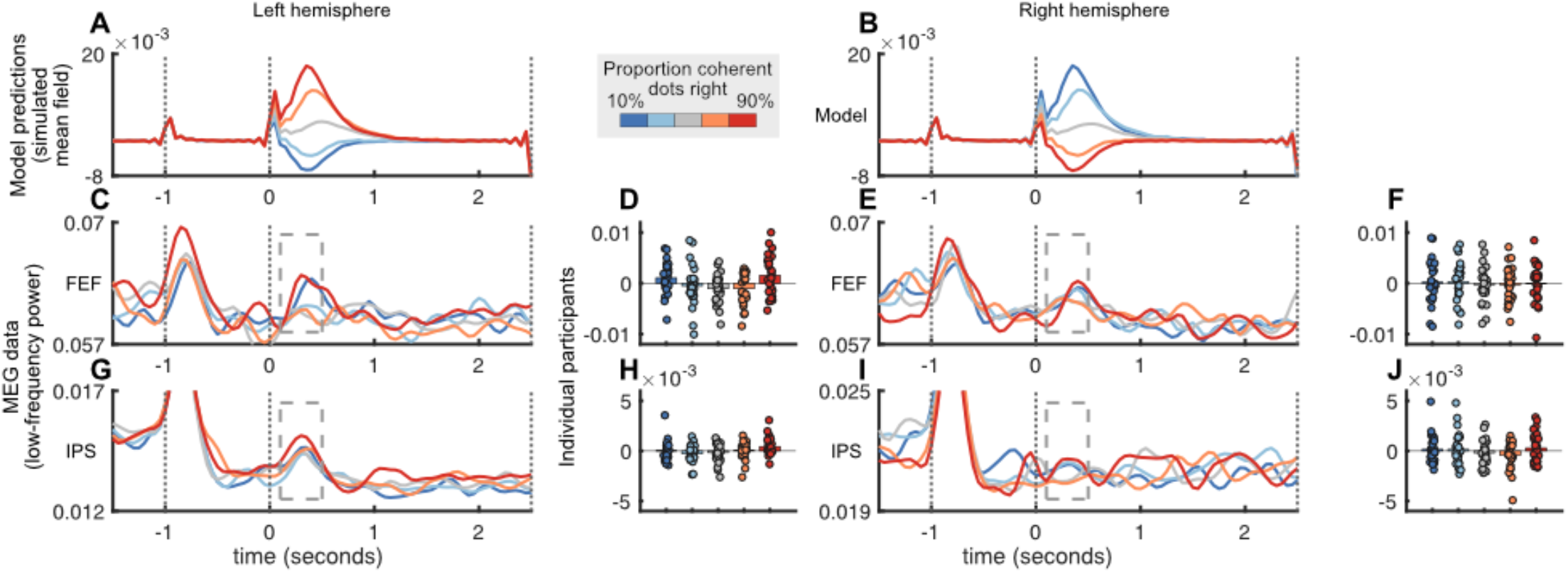
A) Simulated synaptic inputs for right-motion accumulator of neural mean-field model, as a function of percentage coherent dots moving right. B) As A, but left-motion accumulator. C) Source-reconstructed activity from left FEF, same colour convention. D) Individual participant data in stimulus-evoked window. E-F) As C-D, but right FEF. G-J) As C-F but parietal cortex.

Additionally, although we did not detect an overall effect of the prior on evidence accumulation signals in the ERF, for completeness we tested whether a lateralized effect of the prior could be found. To generate predictions we re-ran our neural mean-field model with five levels of pre-stimulus input, ranging from ‘strong left’ (strong input to left motion pool, no input to right motion pool), to ‘strong right’ (reversed). This produced a linear modulation of the simulated evoked response that was proportional to the strength of the pre-stimulus input (figS6A,B).

To compare with our MEG data we fit a general linear model, with prior belief predicted by the Bayesian model as a regressor. However, our prediction was not confirmed (figS6C-F). In contrast to the neural mean-field model, no evidence was observed of linear trends in the MEG data as a function of prior belief (left, t(25) = 1.46, p = 0.16, right, t(25) = −0.52, p = 0.61, figS6C,F). Qualitatively, a quadratic effect was observed whereby weak prior belief produced a strong evoked response and strong belief in either direction produced a weak response. This could reflect an overall process by which a competitive system in FEF is engaged more strongly when the prior is weak; however, this prediction was outside the scope of our model.

**Figure S6:**
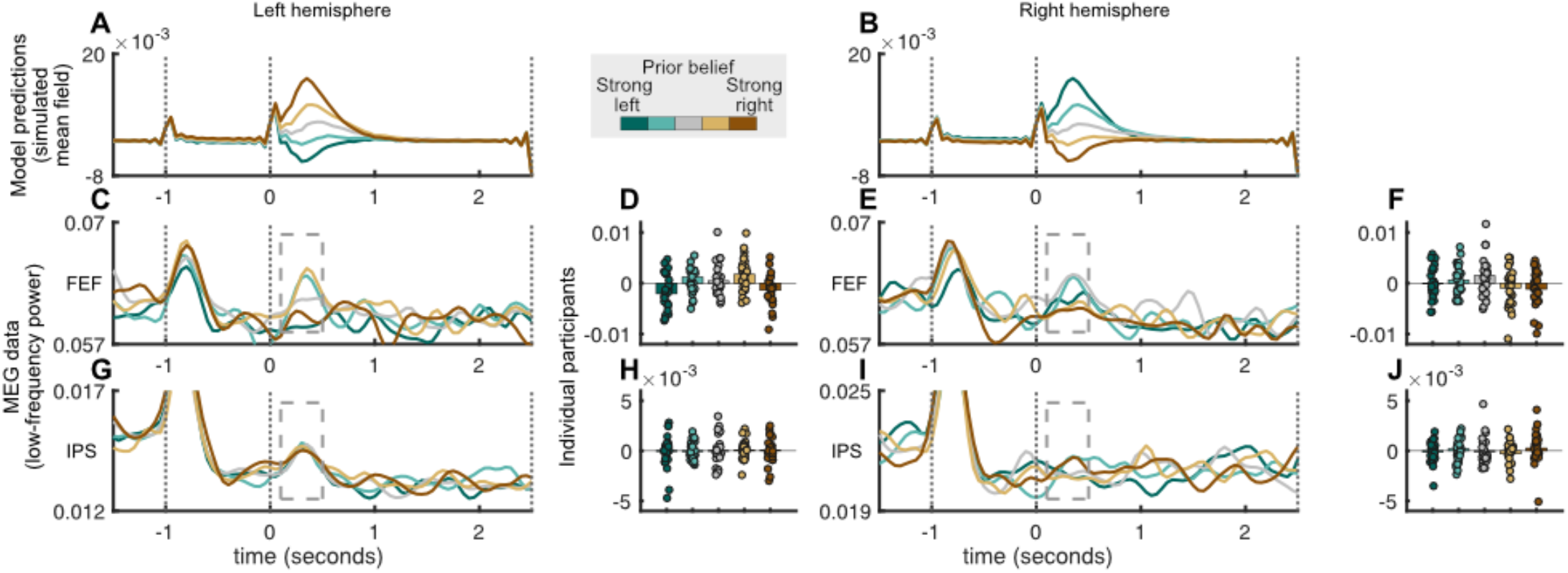
A) Simulated synaptic inputs for right-motion accumulator of neural mean-field model as a function of prior belief. B) Source reconstructed activity in left FEF. C) Individual participant data in stimulus-evoked window. D-F) As A-C, but left-motion accumulator and right FEF.

In parietal cortex we observed a different pattern: Here, no significant linear effect was observed in left parietal cortex (t(25) = 0.88, p = 0.39, figS6G,H). A significant trend was observed in right parietal cortex (t(25) = 2.18, p = 0.039, figS6I,J), but this was in the opposite direction to the prediction from the neural mean field model, with strong *ipsilateral* (rightward) belief showing the largest response.

## References

1. Yarbus, A. L. Eye movements and vision. (1967). doi:10.1016/0028-3932(68)90012-2

2. Land, M., Mennie, N. & Rusted, J. The roles of vision and eye movements in the control of activities of daily living. Perception 28, 1311–1328 (1999).

3. Itti, L. & Baldi, P. Bayesian surprise attracts human attention. Vision Res. 49, 1295–1306 (2009).

4. Itti, L. & Koch, C. Computational modelling of visual attention. Nat. Rev. Neurosci. 2, 194–203 (2001).

5. Usher, M. & McClelland, J. L. The time course of perceptual choice: The leaky, competing accumulator model. Psychol. Rev. 108, 550–592 (2001).

6. Amit, R., Abeles, D., Bar-Gad, I. & Yuval-Greenberg, S. Temporal dynamics of saccades explained by a self-paced process. Sci. Rep. (2017). doi:10.1038/s41598-017-00881-7

7. Gottlieb, J. P. From Thought to Action: The Parietal Cortex as a Bridge between Perception, Action, and Cognition. Neuron 53, 9–16 (2007).

8. Koch, C. & Ullman, S. Shifts in selective visual attention: towards the underlying neural circuitry. Human Neurobiology 4, 219–227 (1985).

9. Friston, K., Thornton, C. & Clark, A. Free-energy minimization and the dark-room problem. Front. Psychol. 3, 1–7 (2012).

10. Kim, J. N. & Shadlen, M. N. Neural correlates of a decision in the dorsolateral prefrontal cortex of the macaque. Nat. Neurosci. 2, 176–185 (1999).

11. Roitman, J. D. & Shadlen, M. N. Response of neurons in the lateral intraparietal area during a combined visual discrimination reaction time task. J. Neurosci. 22, 9475–9489 (2002).

12. Gold, J. I. & Shadlen, M. N. The neural basis of decision making. Annu. Rev. Neurosci. 30, 535–574 (2007).

13. Ding, L. & Gold, J. I. Neural correlates of perceptual decision making before, during, and after decision commitment in monkey frontal eye field. Cereb. Cortex 22, 1052–1067 (2012).

14. Newsome, W. T. & Pare, E. B. A selective impairment of motion perception following lesions of the middle temporal visual area (MT). J. Neurosci. 8, 2201–2211 (1988).

15. Shadlen, M. N. & Newsome, W. T. Neural basis of a perceptual decision in the parietal cortex (area LIP) of the rhesus monkey. J Neurophysiol 86, 1916–1936 (2001).

16. Bogacz, R., Brown, E., Moehlis, J., Holmes, P. & Cohen, J. D. The physics of optimal decision making: A formal analysis of models of performance in two-alternative forced-choice tasks. Psychol. Rev. 113, 700–765 (2006).

17. Soltani, A. & Wang, X. J. Synaptic computation underlying probabilistic inference. Nat. Neurosci. 13, 112–119 (2010).

18. Wong, K.-F. & Wang, X.-J. A Recurrent Network Mechanism of Time Integration in Perceptual Decisions. J. Neurosci. 26, 1314–1328 (2006).

19. Wong, K. F. & Wang, X. J. A recurrent network mechanism of time integration in perceptual decisions. J. Neurosci. (2006). doi:10.1523/JNEUROSCI.3733-05.2006

20. Stokes, M. G. ‘Activity-silent’ working memory in prefrontal cortex: A dynamic coding framework. Trends Cogn. Sci. 19, 394–405 (2015).

21. Drijvers, L., Jensen, O. & Spaak, E. Rapid invisible frequency tagging reveals nonlinear integration of auditory and visual information. Hum. Brain Mapp. 1–15 (2020). doi:10.1002/hbm.25282

22. Zhigalov, A., Herring, J. D., Herpers, J., Bergmann, T. O. & Jensen, O. Probing cortical excitability using rapid frequency tagging. Neuroimage 195, 59–66 (2019).

23. Akrami, A., Kopec, C. D., Diamond, M. E. & Brody, C. D. Posterior parietal cortex represents sensory history and mediates its effects on behaviour. Nature 554, 368–372 (2018).

24. Erlich, J. C., Brunton, B. W., Duan, C. A., Hanks, T. D. & Brody, C. D. Distinct effects of prefrontal and parietal cortex inactivations on an accumulation of evidence task in the rat. Elife 4, 1–28 (2015).

25. Hanks, T. D. et al. Distinct relationships of parietal and prefrontal cortices to evidence accumulation. Nature 520, 220–3 (2015).

26. Britten, K. H., Shadlen, M. N., Newsome, W. T. & Movshon, J. A. The analysis of visual motion: a comparison of neuronal and psychophysical performance. J. Neurosci. 12, 4745–4765 (1992).

27. O’Reilly, J. X., Jbabdi, S., Rushworth, M. F. S. & Behrens, T. E. J. Brain Systems for Probabilistic and Dynamic Prediction: Computational Specificity and Integration. PLoS Biol. 11 (2013).

28. Behrens, T. E. J., Woolrich, M. W., Walton, M. E. & Rushworth, M. F. S. Learning the value of information in an uncertain world. Nat. Neurosci. 10, 1214–21 (2007).

29. Wong, K.-F., Huk, A. C., Shadlen, M. N. & Wang, X.-J. Neural circuit dynamics underlying accumulation of time-varying evidence during perceptual decision making. Front. Comput. Neurosci. 1, 6 (2007).

30. Hunt, L. T. et al. Mechanisms underlying cortical activity during value-guided choice. Nat. Neurosci. 15, 470–476 (2012).

31. Cox, R. T. Probability, Frequency and Reasonable Expectation. Am. J. Phys. (1946). doi:10.1119/1.1990764

32. Bonaiuto, J. J., De Berker, A. & Bestmann, S. Response repetition biases in human perceptual decisions are explained by activity decay in competitive attractor models. Elife (2016). doi:10.7554/eLife.20047

33. Hunt, L. T., Behrens, T. E. J., Hosokawa, T., Wallis, J. D. & Kennerley, S. W. Capturing the temporal evolution of choice across prefrontal cortex. Elife 4, e11945 (2015).

34. Hanks, T. D., Mazurek, M. E., Kiani, R., Hopp, E. & Shadlen, M. N. Elapsed decision time affects the weighting of prior probability in a perceptual decision task. J. Neurosci. 31, 6339–6352 (2011).

35. Platt, M. L. & Glimcher, P. W. Neural correlates of decision variables in parietal cortex. Nature 400, 233–238 (1999).

36. Hauser, C. K., Zhu, D., Stanford, T. R. & Salinas, E. Motor selection dynamics in FEF explain the reaction time variance of saccades to single targets. Elife (2018). doi:10.7554/eLife.33456

37. Van Veen, B. D., van Drongelen, W., Yuchtman, M. & Suzuki, A. Localization of brain electrical activity via linearly constrained minimum variance spatial filtering. IEEE Trans. Biomed. Eng. 44, 867–80 (1997).

38. Hämäläinen, M., Hari, R., Ilmoniemi, R. J., Knuutila, J. & Lounasmaa, O. V. Magnetoencephalography theory, instrumentation, and applications to noninvasive studies of the working human brain. Rev. Mod. Phys. 65, 413–497 (1993).

39. Katz, L. N., Yates, J. L., Pillow, J. W. & Huk, A. C. Dissociated functional significance of decision-related activity in the primate dorsal stream. Nature 535, 285–288 (2016).

40. Adrian, E. D. & Matthews, B. H. C. The Berger Rhythm: Potential Changes from the Occipital Lobes in Man. Brain 57, 355–385 (1934).

41. Müller, M. M., Malinowski, P., Gruber, T. & Hillyard, S. A. Sustained division of theattentional spotlight. Nature 424, 309–312 (2003).

42. Moratti, S., Clementz, B. A., Gao, Y., Ortiz, T. & Keil, A. Neural mechanisms of evoked oscillations: Stability and interaction with transient events. Hum. Brain Mapp. 28, 1318–1333 (2007).

43. Thomas, N. W. D. & Paré, M. Temporal processing of saccade targets in parietal cortex area LIP during visual search. J. Neurophysiol. 97, 942–947 (2007).

44. Colby, C. L., Duhamel, J. R. & Goldberg, M. E. Visual, presaccadic, and cognitive activation of single neurons in monkey lateral intraparietal area. J. Neurophysiol. 76, 2841–2852 (1996).

45. Bisley, J. W. & Goldberg, M. E. Neuronal Activity in the Lateral Intraparietal Area and Spatial Attention. Science (80-.). 299, 81–86 (2003).

46. Ipata, A. E., Gee, A. L., Gottlieb, J. P., Bisley, J. W. & Goldberg, M. E. LIP responses to a popout stimulus are reduced if it is overtly ignored. Nat. Neurosci. 9, 1071–1076 (2006).

47. Mars, R. B. et al. Diffusion-Weighted Imaging Tractography-Based Parcellation of the Human Parietal Cortex and Comparison with Human and Macaque Resting-State Functional Connectivity. J. Neurosci. 31, 4087–4100 (2011).

48. Wimmer, K., Nykamp, D. Q., Constantinidis, C. & Compte, A. Bump attractor dynamics in prefrontal cortex explains behavioral precision in spatial working memory. Nat. Neurosci. 17, 431–439 (2014).

49. Barbosa, J. et al. Interplay between persistent activity and activity-silent dynamics in the prefrontal cortex underlies serial biases in working memory. Nat. Neurosci. 23, 1016–1024 (2020).

50. Seijdel, N., Marshall, T. R. & Drijvers, L. Rapid invisible frequency tagging (RIFT): a promising technique to study neural and cognitive processing using naturalistic paradigms. Cereb. Cortex 1–4 (2022). doi:10.1093/cercor/bhac160

51. Wolff, M. J., Jochim, J., Akyurek, E. G. & Stokes, M. G. Dynamic hidden states underlying working memory guided behaviour. Nat. Neurosci. 1–35 (2017). doi:10.1038/nn.4546

52. Rosner, B. Percentage points for a generalized esd many-outlier procedure. Technometrics 25, 165–172 (1983).

53. Oostenveld, R., Fries, P., Maris, E. & Schoffelen, J.-M. FieldTrip: Open source software for advanced analysis of MEG, EEG, and invasive electrophysiological data. Comput. Intell. Neurosci. 2011, 156869 (2011).

54. Woolrich, M. W., Hunt, L. T., Groves, A. & Barnes, G. R. MEG beamforming using Bayesian PCA for adaptive data covariance matrix regularization. Neuroimage 57, 1466–1479 (2011).

55. Quinn, A. J. et al. Task-evoked dynamic network analysis through Hidden Markov Modeling. Front. Neurosci. 12, 1–17 (2018).

56. Colclough, G. L., Brookes, M. J., Smith, S. M. & Woolrich, M. W. A symmetric multivariate leakage correction for MEG connectomes. Neuroimage 117, 439–448 (2015).

57. Wang, L., Mruczek, R. E., Arcaro, M. J. & Kastner, S. Probabilistic maps of visual topography in human cortex. Cereb. Cortex 25, 3911–3931 (2015).

58. Maris, E. & Oostenveld, R. Nonparametric statistical testing of EEG-and MEG-data. J. Neurosci. Methods 164, 177–90 (2007).

59. Bruce, C. J., Goldberg, M. E., Bushnell, M. C. & Stanton, G. B. Primate frontal eye fields. II. Physiological and anatomical correlates of electrically evoked eye movements. J. Neurophysiol. 54, 714–734 (1985).

60. Bruce, C. J. & Goldberg, M. E. Primate frontal eye fields. I. Single neurons discharging before saccades. J. Neurophysiol. 53, 603–635 (1985).

61. Mesulam, M.-M. Spatial attention and neglect: parietal, frontal and cingulate contributions to the mental representation and attentional targeting of salient extrapersonal events. Philos. Trans. R. Soc. London 354, 1325–1346 (1999).

